# Axonal plasticity in response to active forces generated through magnetic nanopulling

**DOI:** 10.1101/2022.04.12.487762

**Authors:** Alessandro Falconieri, Sara De Vincentiis, Valentina Cappello, Domenica Convertino, Samuele Ghignoli, Sofia Figoli, Stefano Luin, Frederic Català-Castro, Laura Marchetti, Ugo Borello, Michael Krieg, Vittoria Raffa

**Author notes:** equally contributing (Alessandro Falconieri, Sara De Vincentiis).

## Abstract

Mechanical force is crucial in guiding axon outgrowth, before and after synapse formation. This process is referred to as “stretch-growth”. However, how neurons transduce mechanical inputs into signaling pathways remains poorly understood. Another open question is how stretch-growth is coupled in time with the intercalated addition of new mass along the entire axon. Here, we demonstrate that active mechanical force generated by magnetic nano-pulling induces a remodeling of the axonal cytoskeleton. Specifically, the increase in the axonal density of microtubules leads to an accumulation of organelles and signaling vesicles which, in turn, promotes local translation by increasing the probability of assembly of the “translation factories”. The modulation of axonal transport and local translation sustains enhanced axon outgrowth and synapse maturation.

## Introduction

Mechanical force is one of the major extrinsic factors driving axonal outgrowth (Franze, 2013; Suter and Miller, 2011). Neurons generate the force required for motion during axonal elongation and pathfinding. They transduce exogenous forces into signaling, which makes them mechanosensitive cells. In developing neurons, the growth cone (GC) guides axon extension, by detecting guidance cues (Tamariz and Varela-Echavarría, 2015). The study of the molecular mechanisms governing the trip of the tip highlighted an important contribution from signal mechanotransduction (Lowery and Van Vactor, 2009). In GCs, the actin cytoskeleton is connected to the matrix via point contact adhesions. Exogenous pulling forces can trigger the maturation of adhesions and acto-myosin contraction which, in turn, results in the advance of GC through protrusion of actin filopodia/lamellipodia and translocation of microtubules (MTs) in the adhesion site (Lee and Suter, 2008). Interestingly, some guidance cues, such as nerve growth factor (NGF) and netrin-1, appear to activate the same cascade of events upon binding with their own receptors (Toriyama et al., 2013; Turney et al., 2016). However, several lines of evidence suggest that GC is not the only protagonist in signal mechanotransduction in axon outgrowth (de Rooij et al., 2018). First, after connecting with the target, integrated axons continue to grow, thereby accommodating the increase in body mass (Weiss and Hiscoe, 1948). Second, neurites can elongate even if GC and filopodial movements are inhibited (Marsh and Letourneau, 1984). Third, stretched axons elongate at a higher rate than the rate imposed by GC (Pfister et al., 2004), but they are similar to naturally grown axons in terms of structure and ability to transmit electrical signals (Magdesian et al., 2016). The GC even limits the axon to fully exploit its intrinsic capacity to elongate (Lamoureux et al., 1989; Steketee et al., 2014). Importantly, recent studies have contradicted the view that mass is added exclusively at the tip, while the axon shaft remains stationary. Conversely, the intercalated addition of lipids, proteins, vesicles, and organelles along the stretched axon has been demonstrated (Bray, 1984; Lamoureux et al., 2010; Miller and Sheetz, 2006). However, it is not clear which mechanisms underlie the addition of these novel components in the axon. Similarly, although the effects of exogenous forces on the local remodeling of the GC cytoskeleton are well understood, little is known about the axonal plasticity in response to active mechanical stimuli, mainly because of the limited availability of suitable biophysical tools. Magnetic manipulation is now being used to chronically expose axons of developing neurons to extremely low (picoNewton, pN) mechanical forces (Chowdary et al., 2019; Gahl and Kunze, 2018; Kunze et al., 2015, 2017; Raffa et al., 2018; Riggio et al., 2014; Steketee et al., 2014; Tay et al., 2016; de Vincentiis et al., 2020; Wang et al., 2020). Briefly, the protocol is based on the whole axon labeling with magnetic nanoparticles (MNPs) as well as the use of a magnetic field gradient to generate a magnetic force on MNPs tightly interacting with the elastic components (*e.g*. cell membranes or the cytoskeletal network) of the axon. The forces generated by the magnetic nano-pulling appear to modulate axonal guidance (Riggio et al., 2014), elongation and sprouting (Raffa et al., 2018; de Vincentiis et al., 2020), vesicle transport (Chowdary et al., 2019, 2013; Kunze et al., 2017), neuron polarity (Kunze et al., 2015) and differentiation of neural precursor cells (Dai et al., 2019). In our study, we used magnetic nano-pulling to answer some open questions regarding the role of external forces on axon outgrowth. Does the stretch induce axonal cytoskeleton remodeling? Is this related to the addition of novel components? How is this linked to axon outgrowth and synaptogenesis? We focused on the contribution of axonal MTs, on the alteration of the local transcriptome, and on the local mechanisms responsible for mass addition, *i.e*., local transports, local translation, and the cross-talk between them in response to the stimulation.

## Results

### Axonal microtubules sustain stretch-growth

We used a multi-chamber microfluidic device to spatially segregate mouse hippocampal axons from their cell bodies and dendrites (Kim et al., 2012; Taylor et al., 2005). When these axons undergo magnetic nano-pulling, they increase significantly in length (526.9±22.2 μm and 719.4±17.52 μm for control and stretched axons, respectively; the stretching time (*t_s_*) was 120 h, Fig. 1A3, *p*<0.0001), in line with similar data already published by our team (Raffa et al., 2018; de Vincentiis et al., 2020). We then used TEM analysis to understand whether the force generated by the nano-pulling induces a remodeling of the neurite cytoskeleton. Microtubules were found to maintain a normal polarity, morphology and integrity (Fig. 1A). After 120h of stimulation, we found a 36% increase in MT linear density in stretched neurites (from 6.05±0.37 μm^-1^ for control to 8.23±0.44 μm^-1^ for stretched group, Fig. 1A4, *p*=0.0041). So why do MTs respond to force, and how does force influence MT dynamics? Our previous data (de Vincentiis et al., 2020) would suggest that the observed increase in axonal MT density could be a consequence of an increase in MT stabilization. In fact, in the aforementioned work, we found that the Nocodazole (which interferes with MT polymerization) but not Paclitaxel (MT-stabilizing activity) had a differential effect on stretched axons versus control ones, causing a total inhibition of stretch-growth without any effect on normally grown axons. The idea that force stabilizes MTs is not new. It is likely that several mechanisms could directly or indirectly contribute to microtubule stabilization: in *in vitro* assays, MT assembly is directly modulated by the application of forces (Akiyoshi et al., 2010; Hamant et al., 2019) while, *in vivo*, microtubule associate proteins (MAPs) are another possible target that modulates the stability/instability dynamics of MTs (Sánchez-Huertas and Herrera, 2021; Trushko et al., 2013). We thus first tested the hypothesis that a direct perturbation of MT organization disrupts stretch-growth. We used the *C. elegans* model and found, similarly to mouse hippocampal neurons, a significantly increased length of the axons of wildtype touch receptor neurons (Fig. 1B1) or motor-neurons (Fig. S2A) in the stretched condition. We then repeated the assay with the *C. elegans* strain mutated in *mec-12*, encoding for the major α-tubulin in touch receptor neurons, which is then assembled into 15 protofilament MTs together with MEC-7 (Lockhead et al., 2016) (Fukushige et al., 1999). We found no difference in axon length between the control and stretched samples (Fig. 1B1, *p*=0.49), suggesting that the *C. elegans* neurons lacking *α-tubulin* do not respond to the nano-pulling. To corroborate the findings with MEC-12, we used a mutant strain with a *mec-7* allele in touch receptor neurons that lacks stable 15 protofilament MTs but still forms small diameter 11 protofilament MTs (Zheng et al., 2017). We reasoned that MTs are destabilized at higher temperatures (16°C vs 25°C), due to their faster depolymerization rate at 25°C (Li and Moore, 2020). In line with this rationale, the axon length of stretched samples at 16°C was significantly increased compared to the control (unstretched, *p*<0.0001), whereas neurons cultured at 25°C showed no differences between stretched and control conditions (Fig. 1B2, *p*=0.99). This strongly suggests that stretch-growth is dependent on functional β-tubulin. To validate that stretch-growth is not systemically inhibited at 25°C, we cultured wildtype touch receptor neurons at various temperatures. Even though the absolute axon length at lower temperature was shorter, we observed stretch-growth at both 16 and 25°C (Fig. S2B). Together, these data confirm our previous observation that stretch-growth depends on microtubule formation. In addition, we quantified the fluorescence levels of PTL-1, a microtubule-associated protein with tau-like repeats (Goedert et al., 1996), as a marker of MT abundance. We leveraged the *mec-7* allele used before, which carries a mNeonGreen insertion at the endogenous *ptl-1* locus (Krieg et al., 2017), thus avoiding overexpression artifacts. Interestingly, at 16°C, we found a strong increase in the level of PTL-1 total fluorescence (Fig. S2C, *p*<0.0001) and mean fluorescence (Fig. 1B3, *p*<0.0001) in stimulated axons, while at 25°C there were no differences between control and pulled samples (total fluorescence, *p*>0.999, Fig. S2C; mean fluorescence, *p*>0.999, Fig. 1B3). Considering that no differences in the length of control and stretched axons were observed at 25°C, this indicates that the levels of microtubule-associate protein PTL-1 increases during stretch-growth. The increase in the length of stretched axons at 16°C can explain the increase in PTL-1 total fluorescence but not the increase in PTL-1 mean fluorescence, which can be explained by an increase in MT density.

**Figure 1.**
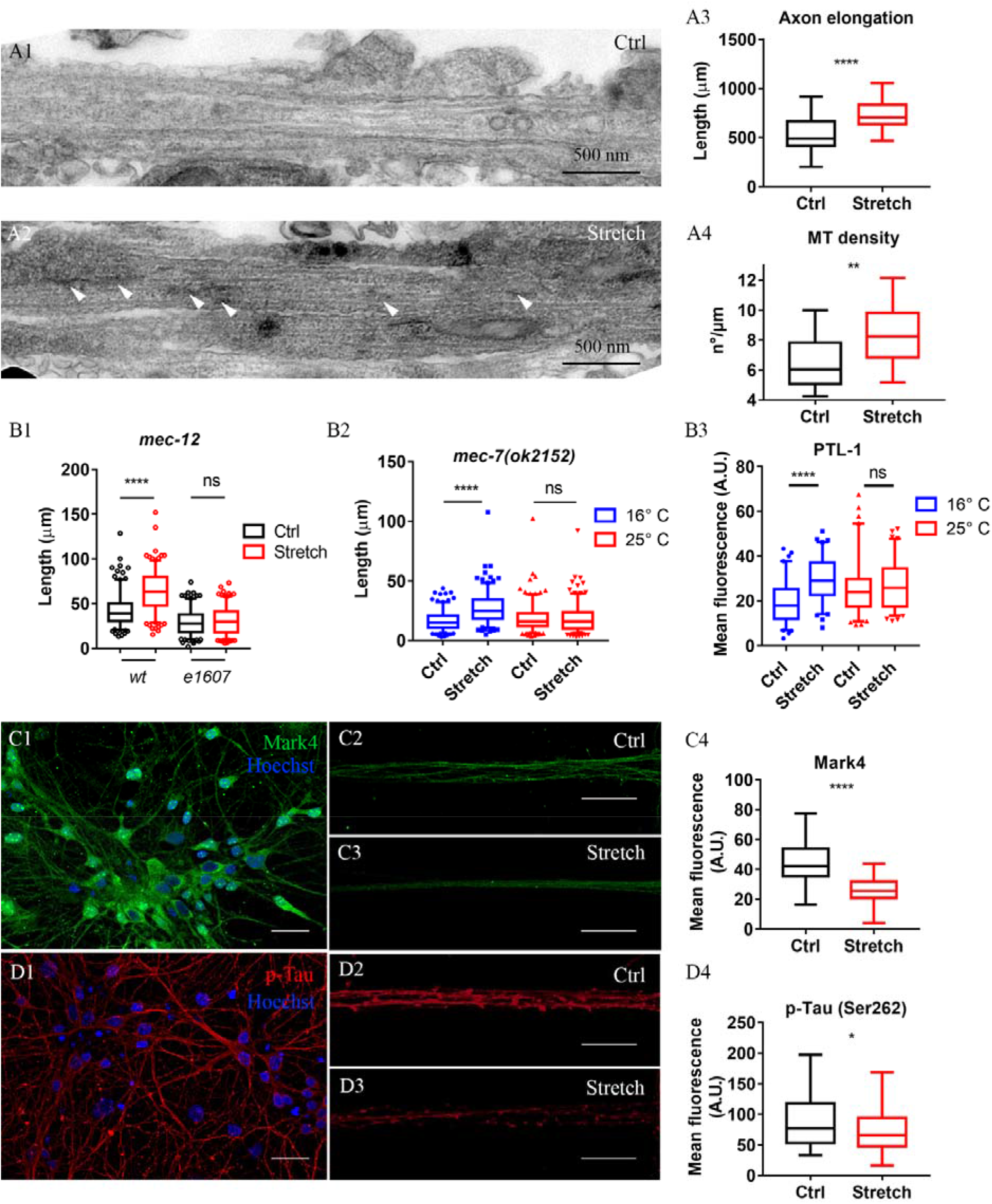
Axon remodeling in response to nano-pulling. A, C, D) DIV6 hippocampal neurons (stretching time, *t_s_*=120 h). B) DIV3 *C. elegans* neurons (*t_s_*=48 h). A) TEM images of cross-sections of control neurites (A1) and stretched ones (A2). Arrowheads indicate ER. A3) Axon lengths in the axon compartment. Box plot (min-to-max), n=60 axons from five biological replicates, unpaired *t* test, 2-tailed, *p*<0.0001, t=6.807, df=118. A4) Quantification of MT linear density, n=21 neurites from two biological replicates. Box plot (min-to-max), unpaired *t* test, 2-tailed, *p*=0.0041, t=3,042 df=40. B1) Axonal length of wt and α-tubulin KO touch receptor neurons (*mec-12(e1607)*) in the control and stretched conditions. Box plot (5-95 percentile), n=200 neurons from four biological replicates. Kruskal-Wallis test with post hoc Dunn’s test, *p*<0.0001. B2) Axonal length of touch receptor neurons in a β-tubulin loss-of-function background (*mec-7(ok2152)*) lacking ultrastable 15 protofilament MTs. Control and stretched axons measured at 16°C, when 11 protofilament MTs are present and at 25°C, when most MTs have been depolymerized. Box plot (5-95 percentile), n=200 axons from four biological replicates, Kruskal-Wallis test with post hoc Dunn’s test, *p*<0.0001. B3) Quantification of mNG mean fluorescence in a transgenic model of tagged mNG::PTL-1 at 16°C and at 25°C. Box plot (5-95 percentile), n=80 axons from four biological replicates, Kruskal-Wallis test with post hoc Dunn’s test, *p*<0.0001. C1) Immunohistochemistry against MARK4 (green) in control axons (C2) and stretched ones (C3). Scale bars: 25 μm (C1), 20 μm (C2-3). C4) Quantification of MARK4 mean fluorescence. Box plot (min-to-max), n=80 microfluidic channels from four biological replicates; unpaired *t* test, 2-tailed, *p*<0.0001, t=10,27 df=163. D1) The specific target of MARK4 kinase, TAU (Ser262) (red), was evaluated in control (D2) and stretched (D3) axons. Scale bars: 25 μm (D1), 20 μm (D2-3). D4) Quantification of TAU (Ser262) mean fluorescence. Box plot (min-to-max), n=80 microfluidic channels from four biological replicates, Mann-Whitney test, *p*<0.0001.

We also tested the possible involvement of MAPs in hippocampal neurons. We focused on TAU for the following reasons: i) TAU phosphorylation influences MT assembly (Kadavath et al., 2015); ii) re-positioning of TAU was already found in response to force (Kunze et al., 2015); iii) we found a down-regulation of the kinase microtubule affinity-regulating kinase 4 (MARK4) that phosphorylates TAU on Ser-262 via Axon-seq. The phosphorylation of TAU in Ser-262 residue considerably reduces its binding to MTs, causing MT destabilization (Biernat et al., 1993; Oba et al., 2020; Sengupta et al., 1998). Conversely, when TAU is dephosphorylated, it may promote MT assembly (Lindwall and Cole, 1984; Liu et al., 2007; Scott et al., 1993). Here, we estimated the levels of MARK4 and TAU (Ser262) in stretched versus control axons by immuno-fluorescence and quantification of the mean fluorescence signal. Both the levels of the kinase and its target were decreased in stimulated axons (Fig. 1C-D; *p*<0.0001). Indeed, a tension-dependent down-regulation of MARK4 could be another mechanism contributing to microtubule stabilization by decreasing the levels of TAU (Ser-262).

### Nano-pulling modulates axonal transport

In the axon, the transport of many organelles and vesicles is linked to MTs and MT stability. We analyzed the distribution of endoplasmic reticulum (ER) that appear as ribosome-free membranous structures with a tubular shape parallel to the plasma membrane of the neurites (Fig. 1A2, white arrowheads). Here, we found a strong accumulation of ER in response to the nano-pulling (Fig. 2B). The ER density was 0.17±0.014 and 0.31±0.018 nm per nm^2^ of neurite area for control and stretched groups at DIV3 (*t_s_*=24h, *p*<0.0001), and this trend continued over the following days (0.17±0.012 nm^-1^ for the control and 0.28±0.082 nm^-1^ for the stretched group at DIV6 corresponding to *t_s_*=120h,*p*<0.0001). At later time points (DIV14, corresponding to *t_s_*=13 days), we found a strong increase in the density of mitochondria in neurites (0.052±0.011 and 0.127±0.024 mitochondria per nm^2^ of neurite area for the control and stretched groups, respectively, *p*=0.04), in response to the high energy demands of stretched neurites (Fig. 2A). We found mitochondria in the 25% and in the 34,3%% of the cross-sections for the control and stretched neurites, respectively, and, limiting the analysis to those cross-sections that were positive for the presence of mitochondria, we still found a similar trend (0.207±0.022 and 0.371±0.051 nm^-2^ for the control and stretched neurites, respectively, *p*=0.002).

**Figure 2.**
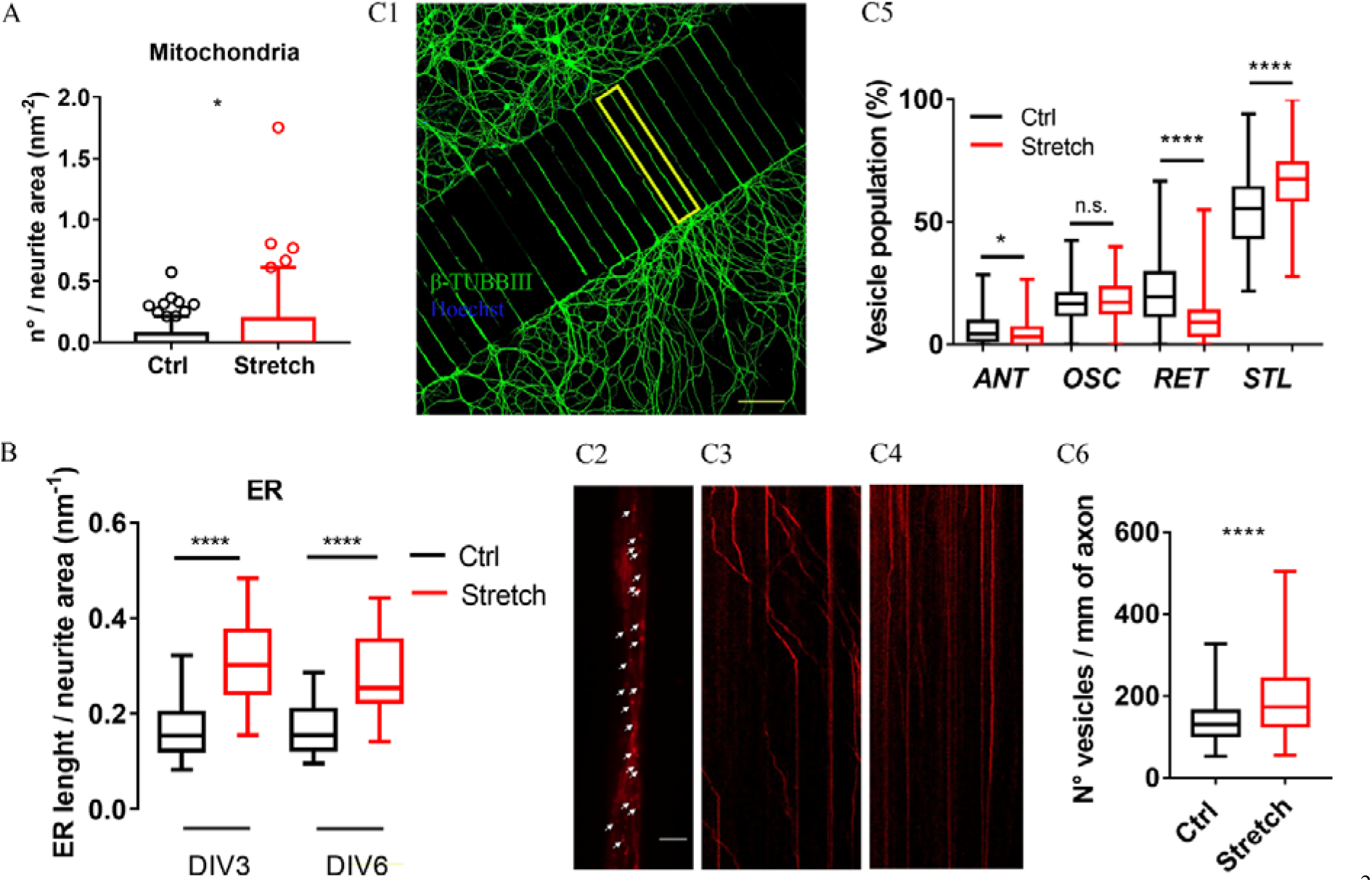
Axonal transport of vesicles and organelles. A) Density of mitochondria (number per nm^2^ of axon area) in control and stretched (*t_s_*=13 days) neurites of DIV14 hippocampal neurons. Box plot (5-95 percentile), n=20 axons from two biological replicates, unpaired *t* test, 2-tailed, *p*=0.0087, t=2,773 df=38. B) Density of ER (nm of tubules per nm^2^ of neurite area) in control and stretched (*t_s_*=2 and 5 days) neurites of DIV3 and DIV6 hippocampal neurons. Box plot (min to max), n=20 neurites from two biological replicates, one-way ANOVA with post hoc Tukey’s comparison, *p*<0.0001, f=22.22. C) Tracking analysis of axonal NGF vesicles in DRG neurons cultured in the microfluidic devices (C1). NGF vesicles (white arrows) in the microfluidic channels (C2, yellow inset of C1). C3-C4) Representative kymographs of the time-dependent displacement of Alexa647-labelled NGF vesicles along a single axon in the control group (C3) and the stretched group (C4); x-scale bar, 10 μm; y-scale bar, 5 s. C5) Total mean population percentages: increase in stalled vesicles and decrease in retrograde ones in stretched groups compared to the controls. Box plot (min to max), three biological replicates, Mann-Whitney test: ANT (*p*=0.01), OSC (*p*=0.12), RET (*p*<0.0001), STL (*p*<0.0001). C6) Density of NGF vesicles in the two conditions: increase in the total number of NGF-vesicles in the stretched group compared to spontaneous elongation. Box plot (min to max), three biological replicates, Mann-Whitney test, *p*<0.0001. β-TUBB3 (green) in (C1); NGF (red) in (C2-4). Scale bars: 50 μm (C1), 5 μm (C2).

Next, we investigated whether the alteration of transport also affects other components, such as signaling vesicles transporting the trophic factors sustaining axon growth (Howe and Mobley, 2004). The NGF was fluorescently labeled, added to the cell growth medium and the axonal trafficking of NGF signaling vesicles was tracked in dorsal root ganglia (DRG) neurons in the presence and absence of the stimulus (Fig. 2C1). To evaluate the impact of forces on the rate of fast axonal transport, we characterized the retrograde (RET), anterograde (ANT), oscillating (OSC), and stalling (STL) components by single vesicle imaging and tracking (Fig. 2C2, inset of C1) (movie S1), as previously reported in (Convertino et al., 2020). Under stimulation, there is a clear accumulation of vesicles in the axon shaft (movie S2). Interestingly, 30 minutes after the magnet removal, axons seem to rescue the levels of control cultures (movie S3). Quantitative data analysis confirmed an overall increase in NGF vesicles in the shaft of stretched axons compared to spontaneous elongation (Fig. 2C6, *p*<0.001). However, the vesicle accumulation is not the consequence of the increase in the anterograde component. In fact, mechanical stimulation led to a strong reduction of retrograde component (*p*=0.01) and to a corresponding increase in stalling ones (*p*<0.0001) (Fig. 2C5). This result was also confirmed by kymograph analysis, which showed more vertical lines in the kymograph of the stretched group, indicating an increased number of stationary vesicles (Fig. 2C4). On the other hand, the kymograph of the control group showed diagonal lines representing moving vesicles (Fig. 2C3). This all suggests a tendency to retrograde vesicles so that they stall in response to the tension.

### Nano-pulling activates local translation

Axonal transport and local translation are the two main mechanisms that sustain the addition of mass in the axon. In particular local translation has an important role in axon elongation and synapse formation (Holt et al., 2019). As a proxy of the rate of axonal translation, we estimated concentration of axonal ribosomes that are in a stage of active translation by immuno-fluorescence (Fig. 3A1) and quantification of the mean fluorescence signal, respectively, in stretched versus unstretched axons. Experimental data revealed a strong increase in the signals of axons undergoing nano-pulling (mean fluorescence, Fig. 3A2, *p*<0.0001). Ribosomes can be assembled in the nucleolus and transported to the axon or assembled locally in the axon in a nucleolus-independent manner (Nagano and Araki, 2021; Nagano et al., 2020; Shigeoka et al., 2019). Whatever the assembly mechanism, axons appear to have control mechanisms to switch on local translation, based on the recruitment of vesicles containing RNA granules, such as late endosomes (Cioni et al., 2019), lysosomes (Liao et al., 2019), and mitochondria (Spillane et al., 2013). Our evidence suggests the ratio of active to total ribosomes increases in stretched axons (Fig. 3A4, *p*<0.0001), but that there is no increase in the number of total axonal ribosomes. In fact, we found a lower concentration of ribosomes in stretched axons, suggesting that the pre-assembled ribosomes and the *in situ* ribosomal components dilute and spread in the longer stretched axons (40% decrease in mean fluorescence signal, from 57.59±3.28 value for the control group to 34.74±2.10 value for the stretch group, Fig. 3B4, *p*<0.0001). This idea is supported by the finding that the concentration of RNA granules shows a similar trend (54% decrease in the volume occupied by RNA granules in the axon, normalized to axon area; *i.e*., 0.057±0.0076 μm and 0.026±0.0042 μm for control and stretched axons, respectively, Fig. 3B3, *p*<0.0001).

**Figure 3.**
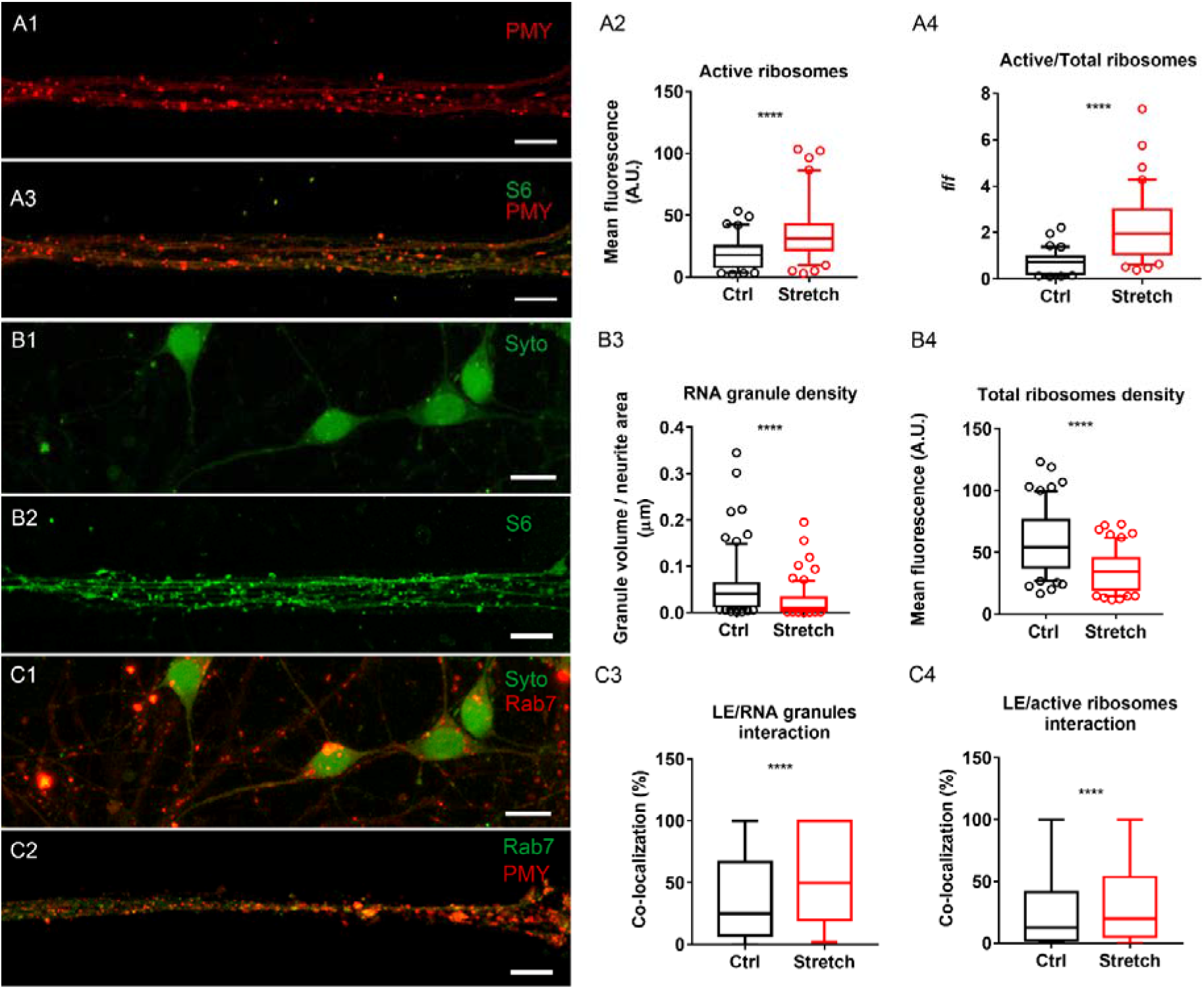
Activation of local translation following nano-pulling in axons of mice hippocampal neurons (DIV6). A1) Staining of ribosome in active translation: PMY (red); scale bars: 10 μm. A2) Increase in mean fluorescence of active ribosomes in stretched versus control axons. Box plot (5-95 percentile), n=80 microfluidic channels from four biological replicates, Mann-Whitney test, *p*<0.0001. A3) Staining of total ribosomes and ribosome in active translation: PMY (red), S6 (green); scale bars: 10 μm. A4) Active ribosomes compared to the total (ratio between fluorescence signals). Box plot (5-95 percentile), n=80 microfluidic channels from three biological replicates, Mann-Whitney test, *p*<0.0001. B1) Staining of RNA granules: Syto RNA select (green); scale bars: 10 μm. B2) Staining of total ribosomes: S6 (green); scale bars: 10 μm. B3) Axonal density of RNA granules expressed as total volume of RNA granules per axon area in control versus stretched axons. Box plot (5-95 percentile), n=75 neurites from three biological replicates, Mann-Whitney test, *p*<0.0001. B4) Quantification of mean fluorescence in control versus stretched axons. Box plot (5-95 percentile), n=60 microfluidic channels from three biological replicates, Mann-Whitney test, *p*<0.0001. C1) Immunostaining of late endosomes (Rab7, red) and RNA granules (Syto RNA select, green); scale bars: 10 μm. C2) Immunostaining of late endosomes (Rab7, green) and active ribosomes (PMY, red) in axons; scale bars: 10 μm. C3) Co-localization between LEs and RNA granules in control versus stretched axons. Box plot (min to max), n=75 neurites from three biological replicates, Mann-Whitney test, *p*<0.0001. C4) Co-localization between LEs and active ribosomes in control versus stretched axons. Box plot (min to max), n=53 microfluidic channels from three biological replicates, Mann-Whitney test, *p*<0.0001.

These data suggest that the stimulus triggers local translation by switching the resident ribosomes from an inactive to active state, rather than promoting their assembly or transport. To support this hypothesis, we analyzed the population of late endosomes. Cioni and colleagues demonstrated that the translation of proteins in axons takes place through intimate contact between late endosomes (LEs) and RNA granules, with LEs becoming the platform for the newly synthesized proteins (Cioni et al., 2019). To establish whether nano-pulling could have an impact on the interaction of these components, we evaluated the co-localizations between LEs and both RNA granules and active ribosomes (Fig. 3C2). Experimental data showed a strong increase in the number of functional interactions between LEs and RNA granules (Fig. 3C3, *p*<0.0001) and between LEs and active ribosomes (Fig. 3C4, *p*<0.0001). Although LEs play different roles in the cell, these data suggest that many of these structures can be engaged in functional interactions for local translation in response to the stimulation. Late endosomes are transported along the MTs tracks through molecular motors (Welte, 2004) and the link between the dynamics of axonal transports and MTs is very strong (Yogev et al., 2016). We speculate that the increase in mitochondria and vesicle density in the axon shaft during the nano-pulling, increases the probability of the formation of functional platforms for local translation.

### Nano-pulling stimulates synapse remodeling

Data we previously collected by electrophysiological recordings showed an enhancement of spontaneous activity and induced currents in stretched axons, both at DIV7 and DIV14 (de Vincentiis et al., 2020). In the light of the data reported in the previous section, we were wondering whether this acceleration of synaptic maturation might be related to an accumulation of synaptic vesicles (SV) and synaptic remodeling. We studied the localization of synapsin I, a protein that is associated with SVs in the synapses of the central nervous system (CNS) (Gitler et al., 2004), as a marker of the initial steps of synapse formation in the developing axon. As the formation of mature synapses comprises several stages, the analysis of synapsin I was carried out on hippocampal neurons which were stimulated for 6, 13, and 20 days of stretching, corresponding to a temporal window spanning from 1 week *in vitro* (DIV6, no synapses) to 3 weeks *in vitro* (DIV21, presence of mature synapses), with 2 weeks as the intermediate stage (DIV 14, immature synapses) as reported for normally grown axons (Muller et al., 1993). According to current understanding, in the developing axons, pre-synaptic proteins are transported in the form of precursor vesicles from the neuronal soma, where they are produced, along axonal MTs to the synaptic terminal (Guedes-Dias and Holzbaur, 2019). During their maturation, there is an extensive remodeling caused by motility, fusion, aggregation, dissociation, “stop and go” trafficking that spans through multiple pre-synaptic sites, clustering, and recycling (Maeder et al., 2014). To take into account this complexity, we analyzed the synapsin I vesicles that appear as fluorescent spots, estimating parameters such as the spot volume (Fig. 4B), spot density (Fig. 4C), and spot fluorescence (Fig. 4D) in control and stretched neurites. These give a rough estimation of the vesicle size, vesicle concentration in the neurites, and synapsin I concentration in the vesicle, respectively. For control cultures, we observed in the experiment timeframe that spots remain constant in volume (Fig. 4B, *p*=0.97), but the fluorescence emitted (Fig. 5D, *p*=0.03), as well as the density (Fig. 4C, *p*=0.02) of the spots, increase at DIV21, corresponding to the time of formation of mature synapses. A multiple comparison among all timepoints highlights that stimulated neurites always have more spots, irrespectively of the time (*p*=0.007). Interestingly, the spots of stimulated neurites are smaller than the control ones at DIV6 (*p*=0.0003), however the significance of this difference is totally lost at DIV21 (*p*=0.56). Although the collected data are insufficient to support any statement, an open question to address in future works could be whether the increase in the linear density of MTs promotes the intense and rapid trafficking of small precursor vesicles. In this respect, it has been suggested (Ahmed and Saif, 2015) that smaller (<350 nm) but not larger (>350 nm) vesicles spend more time undergoing active transport when a neuron is stretched, probably because small vesicles experience less resistance to motion. Overall, our data show that, at DIV21, synapsin I positive spots in stimulated neurites are similar in volume and protein-rich but more numerous than the control ones.

**Figure 4.**
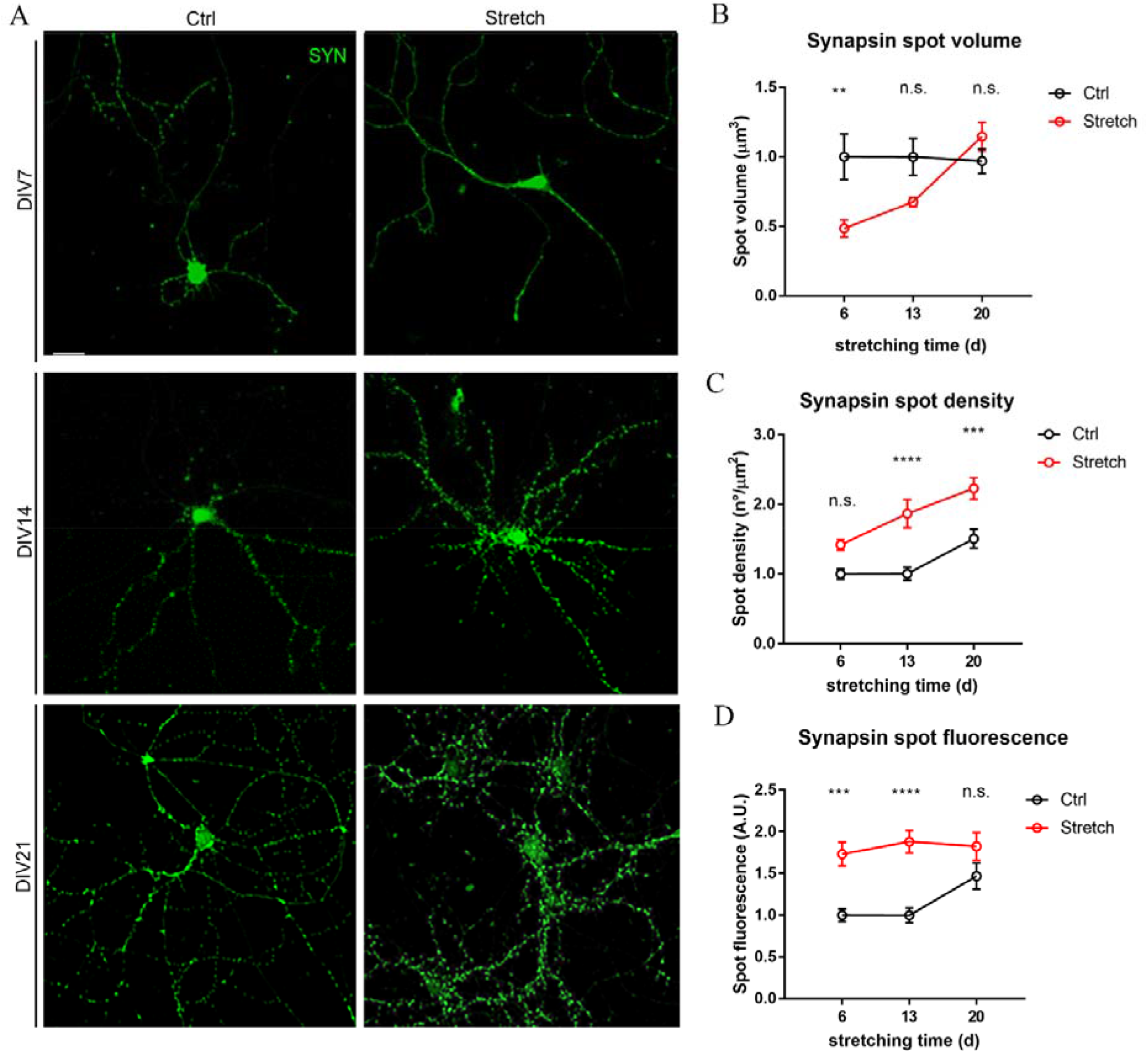
Axonal localization of synapsin I. A) Synapsin I staining (green) of control and stretched neurons at different stretching times. Scale bar: 20 μm. B) Analysis of synapsin I spot volume, C) spot density, and D) spot fluorescence after 6, 13, and 20 days of stimulation (corresponding to DIV7, DIV14, and DIV21, respectively) in control and stretched neurites. For all the panels: Mean ± SEM, n=30 neurons from three biological replicates. Two-way ANOVA with post hoc Sidak’s test. B) Row factor (Ctrl vs Stretch): *p*=0.01, f=4.63. Column factor (days of stretching): *p*=0.01, f=6.47. C) Row factor (Ctrl vs Stretch): *p*=0.007, f=5.03. Column factor (days of stretching): *p*<0.0001, f=40.47. D) Row factor (Ctrl vs Stretch): *p*=0.25, f=1.38. Column factor (days of stretching): *p*<0.0001, f=38.17.

**Figure 5.**
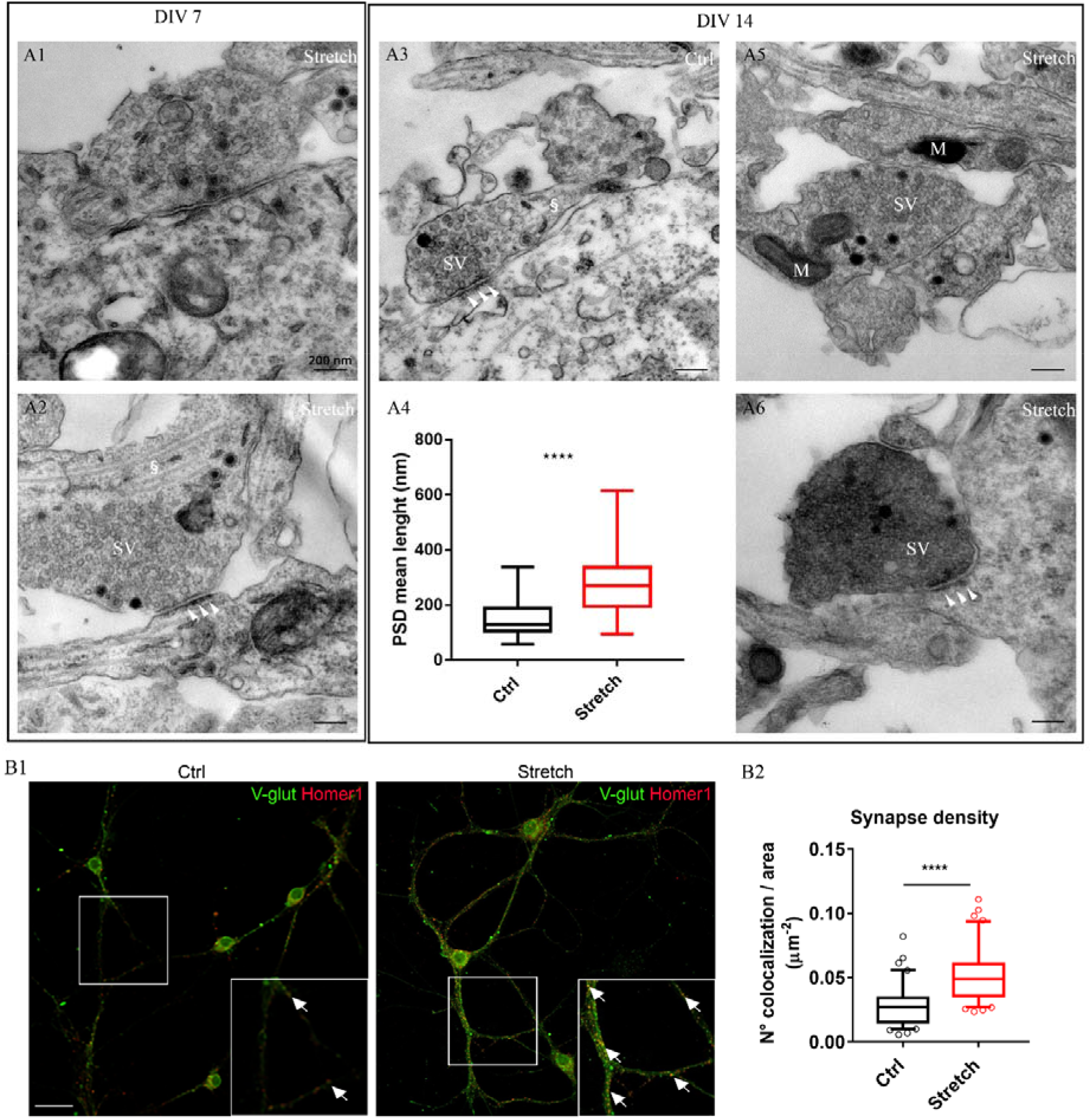
Synapse remodeling. A) Micrographs of synapse “draft” at DIV7 (A1, A2) and DIV14 (A3, A5, A6), in control (A3) and stretched (A1, A2, A5, A6: *t_s_*=13 days) axons of hippocampal neurons. White arrowheads highlight examples of PSD regions, white § MTs, “M” mitochondrion. A4) Length of PSD region in control and stretched (*t_s_*=13 days) axons of DIV14 hippocampal neurons. Box plot (min to max), n>28 axons from two biological replicates, Mann-Whitney test, *p*<0.0001. B1) Immunostaining of V-glut1 (green) and Homer1 (red) in hippocampal neurons at DIV14 in the control and stretched conditions (*t_s_*=13 days). White arrows indicate the co-localizating spots. Scale bar: 20 μm. B2) Quantification of synapse density as the number of colocalizing spots in the selected neurite area in the control and stretched groups. Box plot (10-90 percentile), n=40 neurons from four biological replicates, Mann-Whitney test, *p*<0.0001.

To understand whether the stimulation could speed up synapse formation, the developing axons were analyzed at DIV7 and DIV14 for the presence of SVs, active zone (AZ), synaptic cleft (SC), pre- and post-synaptic membranes, as markers of the presence of a synapse (Liu et al., 2019). At DIV7, no hallmarks of synapses were observed in control conditions (not shown). Conversely, DIV7 neurons stimulated for six days were found to have some developing synapses characterized by the typical pre- and post-synaptic elements on the two sides of the SC. On the presynaptic side we observed the accumulation of SV pools at different distances from the AZ, while, on the post-synaptic side we noted the formation of post-synaptic density (PSD - the specialized region for neurotransmitter recording (Fig. 5A1-A2)). The structure of these synapses was very similar to the ones observed in the control cultures at DIV14 (panels A1-2 versus A3). At DIV14, some synapses of stretched neurons (roughly 15%) were structurally similar to the control ones, in terms of SV density and the presence of MTs (Fig. 5A3 versus A5). However, approximately 85% appeared to be in a more advanced stage, showing a terminal localization and the curved shape typical of mature synapses. In these mature synapses, we also found some in which the presynaptic terminals appeared to be morphologically different from all the others due to a more contrasted cytoplasm (Fig. 5A6 versus A5). This is probably due to a higher density of SVs and also of the actin filaments to which they are anchored, since these are both hallmark features of synapse stabilization and maturation (Zhang and Benson, 2001). In line with this, we also found an increase in the length of PSD (147.9±12.84 nm and 287±19.78 nm for the control and stretch groups, respectively, *p*<0.0001, Fig. 5A4). We also noted an increased number of synapses that show a well-formed PSD (72% in the control group, 95% in the stretched one). We found no spine specialization at these stages.

To get more evidence, the presence of pre- and post-synaptic proteins in close proximity was used as a well-accepted criterion for the identification of a functional synapse (Gürth et al., 2020). Specifically, we measured the co-localization between the presynaptic marker V-glut and the post-synaptic marker Homer1 in neurites at DIV14. There was a significantly higher number of colocalization spots in the stimulated samples than in the control ones (Fig. 5B, p<0.0001). The coupling probability based on the analysis of the number of co-localizing spots normalized for the neurite area considered was determined for the two conditions. The number of co-localizations between pre- and post-synaptic markers was 76±12% higher in the stimulated neurites than in the controls.

### Nano-pulling modifies the axonal transcriptome

To gain an overall perspective of the changes induced by the nano-pulling in the axonal RNAs, we performed the Axon-Seq analysis. RNA was isolated from the somato-dendritic and axonal compartments from DIV6 hippocampal neurons in the control and stretched (*t_s_*=120 h) conditions (n=6). The principal component analysis (PCA) showed that the two compartments have distinct transcriptional profiles (Fig. 6A). We found 907 differentially expressed genes (padj ≤ 0.01 and −2 ≤ log2FoldChange ≤ 2) in axons, and gene ontology (GO) was carried out to investigate which cellular processes were mis-regulated between the two conditions. We used DAVID, a web-based tool, to classify the differentially expressed genes based on the cellular components that they are related to (Huang et al., 2009). Using these criteria, we found many functional categories which had more than 10 transcripts identified as relevant for axons or related processes, *i.e*., organelles (GO:0005794, Golgi apparatus, gene count: 74; GO:0005739, mitochondrion, gene count: 91; GO:0005783, endoplasmic reticulum, gene count: 67), vesicles (GO:0005764, lysosome, gene count: 26; GO:0005765, lysosomal membrane, gene count: 24; GO:0005768, endosome, gene count: 39; GO:0005770, late endosome, gene count: 11; GO:0031902; late endosome membrane, gene count: 12), membrane (GO:0016020, membrane, gene count: 314; GO:0031225, anchored component of membrane, gene count: 13), cytoskeleton organization (GO:0005856, cytoskeleton, gene count: 61; GO:0005815, microtubule organizing center, gene count: 14) and synapse (GO:0045202, synapse, gene count: 28). The GO categories identified (Fig. 6B) reveal many cellular processes that can modulate vesicle trafficking, local transport, local translation machinery, and high energy metabolism, reflecting the need for mass addition and energy demands required to sustain axon outgrowth. Notably, the presence of the categories of cytoskeleton and synapse remodeling reflect the potential role of the stimulus in the enhancement of neuronal maturation. In order to deepen our knowledge on this point, we analysed the 61 dysregulated genes related to cytoskeleton (GO:0005856). In line with the evidence of alterations in the MT density and axonal transport, we found that many upregulated genes are related to MT cytoskeleton organization, MT-binding protein, and MT motor proteins (Fig. 6C, Table S3).

**Figure 6.**
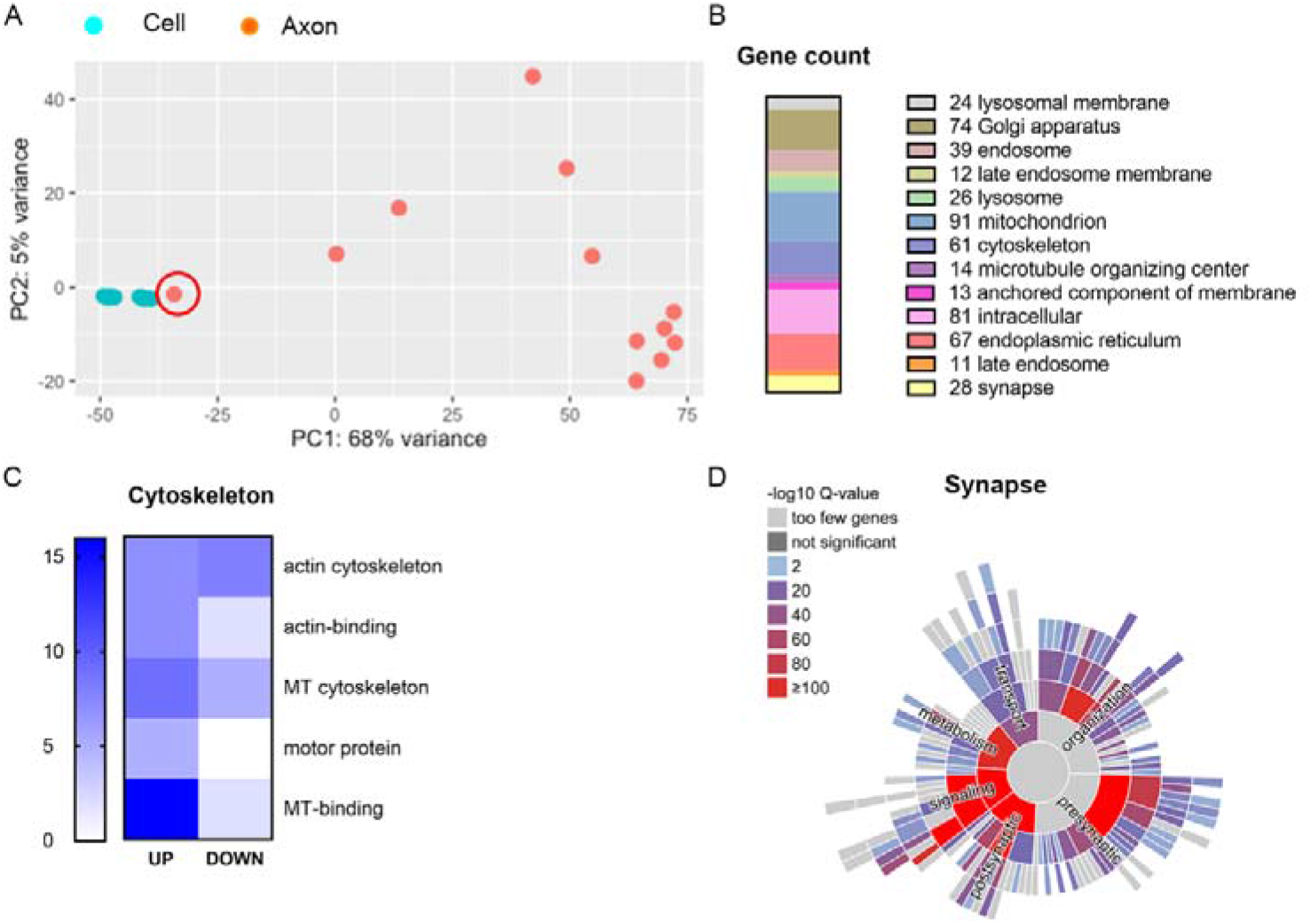
Axon-Seq of hippocampal neurons, *t_s_*: 120 h (from DIV1 to DIV6). A) Principal component analysis. The circled sample highlights an axonal sample contaminated with the somatic component that was excluded from further analysis (see STAR methods). B) Gene ontology via DAVID software, functional annotation chart, GOterm_cc_direct. The graph plots only the axonal or axon-related categories or categories with less than 10 annotated genes. Some general categories (GO:0005737, cytoplasm; GO:0016020, membrane; GO:0005829, cytosol) have not been included but the full list is given in supplementary Table S2. C) Cytoskeleton GO analysis (GO:0005738), manual annotation. The bar plots the gene number. Gene table is provided in supplementary Table S3. D) Synaptic GO analysis via SynGO, function domain, enrichment analysis.

The SynGO tool was used to carry out the same analysis at the synaptic level (Koopmans et al., 2019). There were many genes involved in pre- and post-synaptic dynamics that were found to be differentially expressed in the stretched axons than in the controls. In the first level of the gene ontology enrichment analysis (GOEA), the first-level categories with ≥ 100 unique genes were: synapse organization (GO:0050808, 306 annotated genes), process in the pre-synapse (269 annotated genes), process in the post-synapse (218 annotated genes), synaptic signaling (GO:0099536, 193 annotated genes); the second-level categories with ≥ 50 unique genes were: synapse assembly (GO:0007416, 93 annotated genes), post-synapse organization (GO:0099173, 71 annotated genes), synaptic vesicle cycle (GO:0099504, 73 annotated genes), regulation of post-synaptic membrane neurotransmitter receptor levels (GO:0099072, 121 annotated genes), regulation of post-synaptic membrane potential (GO:0060078, 55 annotated genes), trans-synaptic signaling (GO:0099537, 185 annotated genes) (Fig. 6D).

## Discussion

It is well accepted that neurons are under mechanical tension *in vivo*, and that they shape neuronal morphology and connectivity (Gangatharan et al., 2018)(Krieg et al., 2014). During development and axon outgrowth the GC pulls the axon shaft and this traction force is responsible for intercalated mass addition (Miller and Suter, 2018). The axon continues to be under traction also when the synapse between the axon and its target is established. In fact, the growth of the organism increases the distance between the soma and synapse. The mechanical force is an extrinsic signal that induces axon outgrowth, however the molecular pathways of mechanotransduction are still poorly understood in neurons. The tension generated onto axons by magnetic nano-pulling generates an extremely low force and a constant loading (Chowdary et al., 2013; Dai et al., 2019; Gahl and Kunze, 2018; Kunze et al., 2015; Riggio et al., 2014; Steketee et al., 2014; Tay et al., 2016; de Vincentiis et al., 2020; Wang et al., 2020), similarly to those exerted, during some phases of development, by tissue morphogenesis or, after development, by the post-synaptic muscle tissue. Another novelty of our work is that we used a microfluidic chamber model to isolate the axonal compartment from the somato-dendritic one and to focus on the axonal level. We used these technologies to study axonal remodeling in response to this stimulation. We identified axonal MTs as a promising candidate for orchestrating axon outgrowth and intercalated mass addition. We found an increase in the linear density of MTs in stretched neurites. This difference is maintained during stretching (neurite stretching for 48 or 120 h had a similar increase in MT density compared to the control, *i.e*. about 36%, Fig. 1A4 and (de Vincentiis et al., 2020)). This is not an artifact due to the “inside” action of MNP-mediated nano-pulling as neurites stretched from the “outside” show a similar ultra-structure, in terms of axonal MT accumulation (Falconieri et al., 2021). The crucial role of MTs as a component of the signal mechanotransduction was also corroborated by the analysis of the axonal transcriptome that revealed the many genes related to MT structure, MT-binding proteins and MT axonal transport which were differentially expressed between the stretched and the unstretched conditions (Fig. 6C, Table S3). It is not still clear how the mechanical force can induce an accumulation of MTs in neurites. It is possible that the MTs are mechanosensitive in themselves, since the direct application of force induces their stabilization (Hamant et al., 2019). In *in vitro* assays, MT bundles are able to self-organize and align in the direction of stretch (Inoue et al., 2016). The effect of tension on MTs is direct and an increase in traction force can slow down MT depolymerization or disassembly (Akiyoshi et al., 2010; Hamant et al., 2019). In mammalian cells, the increase in tensile force generated at the adhesion point was found to promote MT assembly (Putnam et al., 2001). Alternatively, an indirect stabilization of MTs can be mediated by MAPs that modulate the stability/instability dynamics of MTs (Sánchez-Huertas and Herrera, 2021), especially in the light of some evidence demonstrating that physical forces can regulate the activity of MAPs. Specifically, a tensile force applied to proteins that bind to the plus-ends of MTs such as XMAP215 in *Xenopus* (Trushko et al., 2013) and Dam1 in yeast (Franck et al., 2007) was found to increase MT length. In this regard, we formulated the data-driven (RNAseq) hypothesis that low MARK4 levels correlated with TAU (ser262) hypo-phosphorylation could also contribute to MT stabilization (Fig. 1C4-D4), however it remains unclear whether the downregulation of MARK4 is mediated by a mechanosensitive protein or a calcium-dependent protein or other mechanisms. A very interesting open issue is how a force of a few pN generated by the nano-pulling can modulate the net contractile force resulting from the pulling forces generated by the GC and the axon shaft which is hundreds of pN (O’Toole et al., 2015). Here, we propose a mechanism defined as “the axonal positive loop” according to which the application of the force of few pN to the axon triggers the accumulation of MTs, each of them generating a force in the range of 4 pN (Kolomeisky and Fisher, 2001), and together they can decrease the pulling force generated by the axon onto the GC by hundreds of pN, thus pushing the tip forward. This mechanism is consistent with the observation that stretch-growth occurs when axons are stretched from the soma to the tip but not from the tip to the soma (Raffa et al., 2018). This is due to the polarity of MTs, which in stage 3 axons, have a preferential polarity with the plus-end facing the tip (Yogev and Shen, 2017). More evidence of the connection between MT stabilization and axon elongation is that the impairment of MT polymerization, via Nocodazole administration, was found to totally block stretch-growth (de Vincentiis et al., 2020), probably by limiting the addition of new mass. In fact, when knocking-out α-tubulin or β-tubulin in touch receptor neurons of *C. elegans*, we found that stretch-growth is not permitted (Fig. 1B). Stretch-growth appears to be well conserved across evolution and strictly dependent on MT dynamics.

One of the most intriguing questions is how the stabilization of MTs can be linked with the role of mechanical tension in mass addition. The GO analysis revealed a functional enrichment in some cellular components such as mitochondria (GO:0005739, gene count 91) and endoplasmic reticulum (GO:0005783, gene count: 67) (Fig. 6B). This may reflect the high energy demands, the lipid and protein hyper-production required to sustain stretch-growth. The data collected in this study thus provide evidence of the accumulation of mitochondria and ER in stretched neurites (Fig. 2A-B). The increase in MT density in the stretched group could account for the accumulation of mitochondria and ER cisternae, whose transport is mostly dependent on microtubule-based motors (Lin and Sheng, 2015),(Farías et al., 2019). MT stability was also found to influence the distribution and structural organization of axonal ER (Farías et al., 2019).

However, the most important result from GOEA is related to vesicle processes, such as lysosome (GO:0005764, gene count: 26), endosome (GO:0005768, gene count: 39), late endosome (GO:0005770, gene count: 11), and synaptic vesicle cycle (GO:0099504, 73 annotated genes). By investigating some vesicular components, we found that nano-pulling promotes an accumulation of synaptic vesicles (Fig. 4) and signaling vesicles (Fig. 2C6). Accumulation of organelles and vesicles could result from the accumulation of MTs given that many of these components have an active motion on MTs (Sánchez-Huertas and Herrera, 2021). However, another mechanism that needs to be considered is that force has been proved to modulate the unbinding rate of molecular motors from MTs (Berger et al., 2019). According to the “tug-of-war” model, a differential response of the kinesin and dynein molecular motors to the force could account for the modulation of the vesicle transport (Leidel et al., 2012). Interestingly, an opposing force of a few pN causes the dynein to walk backwards toward the MT plus end (Gennerich et al., 2007). It would be interesting in future works to investigate whether this force dependent behavior could explain our observation that the accumulation of NGF-vesicles is related to a decrease in the retrograde component rather than to an increase in the anterograde one. Another possibility is that magnetic nano-pulling impairs the retrograde movement of MNP-labeled vesicles (Steketee et al., 2011), however we do not have evidence of the accumulation of MNPs within vesicles.

The model that emerges from this study is that nano-pulling can stimulate the accumulation in the axon of some components of the “translation platforms”. When mitochondria accumulate in the axon shaft, they have a higher probability of associating with RNA granules and of activating local translation (Cioni et al., 2019). In addition, ER tubules that accumulate in stretched axons may also interact with many elements of the translational platforms such as endosomes and mitochondria in order to co-regulate this process (Merianda et al., 2009). The increase in actively-translating ribosomes (Fig. 3A2-A4), but not total ones (Fig. 3B4), together with the increase in the functional interactions between LE and RNA granules (Fig. 3C3) and between LE and active ribosomes (Fig. 3C4) highlight that nano-pulling promotes local translation. These results support the view that the activation of local translation in axons is necessary for stretch-growth. This is also consistent with the fact that the inhibition of protein synthesis via cycloheximide treatment led to the inhibition of stretch-growth but not tip-growth (de Vincentiis et al., 2020).

Our data support the idea that the application of active forces on axons promotes the creation of a complex dialogue between axonal transport and local translation and that this dialogue may be driven by the MTs. This cross-talk could be responsible for the supply of vesicles, lipids and proteins required for sustaining axon outgrowth (Fig. 1A3) and synaptic maturation (Fig. 4, Fig. 5). Further studies are still needed to elucidate the complete signal cascade.

## Acknowledgments

The study was supported by the Wings for Life Foundation (WFL-IT-16/17 and 20/21), the Italian Ministry of Economical Development through the MAECI (MagNerv), the ERC (MechanoSystems, 715243), the HFSP (CDA00023/2018, RGP0026/2021), the Spanish Ministry of Economy and Competitiveness through the Plan Nacional (PGC2018-097882-A-I00), the “Severo Ochoa” program for Centres of Excellence in R\&D (CEX2019-000910-S; RYC-2016-21062), the Fundació Privada Cellex, Fundació Mir-Puig, and the Generalitat de Catalunya through the CERCA and Research program (2017 SGR 1012). We would like to thank the NMSB lab for discussions and Nawaphat Malaiwong in particular for help with *C. elegans* cell culture. The authors would like to thank Alessandra Galeotti for re-analysing the transcriptomic data, and Antonino Cattaneo and Antonietta Calvello of the Bio@SNS lab for making NGF-YBBR available for these studies.

## Author contributions

Conceptualization, M.K., V.R.; Methodology, A.F., V.C., L.M., S.L., F.C.; Formal analysis: U.B.; Investigation, A.F., S.D.V., V.C., D.C., S.G., S.F.; Writing – Original Draft, V.R. and A.F.; Review & Editing, All; Funding Acquisition, M.K. and V.R.; Resources, V.C., S. N., L.M., M.K. and V.R.; Supervision, L.M., U.B., M.K. and V.R.

## Declaration of interests

The authors declare no competing interests.

## STAR methods

### Resource availability

#### Lead contact

All images, data and metadata are stored in Google Drive and are accessible to lab users (https://drive.google.com/drive/folders/1MZuFRWTNnGn7WeSdfFp20wSe_hH0A_Xg). Further information and requests for accessing these resources and reagents should be directed to Vittoria Raffa.

### Materials availability

This study did not generate new unique reagents.

### Experimental model and subject details

#### Mice

Animal procedures were performed in strict compliance with protocols approved by the Italian Ministry of Public Health and the local Ethical Committee of the University of Pisa (protocol number 39E1C.N.5Q7) in conformity with the Directive 2010/63/EU. C57BL/6J mice were kept in a regulated environment (23 ± 1°C, 50 ± 5% humidity) with a 12 hour light-dark cycle with food and water *ad libitum*.

#### C. elegans

We used the following *C. elegans* strains: GN692 [*ljSi123*[*mec-7*p:GCaMP6s::SL2::tagRFP];*lite-1(ce314)*] worm strain, which expresses GCaMP6s and the calcium-independent tagRFP in touch receptor neurons; MSB32 [*hpIs258; lite-1(ce314)* X] which expresses the green calcium sensitive fluorophore in motor-neurons (Wen et al., 2012); GN510 [*mec-4(zdIs5)* I; *mec-12(e1607)* III] worm strain that expresses GCaMP6s and the mutation (*mec-12(e1607)*) for α-tubulin in touch receptor neurons; GN647 [*ptl-1(pg73)* III; *mec-7(ok2152)* X] (Krieg et al., 2017), worm strain mutant for MEC7 β-tubulin in touch receptor neurons lacking stable 15 protofilament microtubules.

### Method details

#### Cell culture

For hippocampal neurons, newborn animals (P1 stage) were sacrificed and both hippocampi were harvested in a solution of D-glucose 6.5 mg·ml^-1^ in DPBS (Gibco, Thermo Fisher Scientific, Waltham, Massachusetts, US, #14190-094). Cell isolation was performed by chemical digestion and mechanical dissociation as previously described (de Vincentiis et al., 2020). After this step, cells were seeded in high-glucose DMEM (Gibco, Thermo Fisher Scientific, Waltham, Massachusetts, US, #21063-029) with 10% fetal bovine serum (FBS, Gibco; Thermo Fisher Scientific, Waltham, Massachusetts, US, #10270-106), 100 IU·ml^-1^ penicillin, 100 μg·ml^-1^ streptomycin (Gibco, Thermo Fisher Scientific, Waltham, Massachusetts, US, #15140-122) and 2 mM Glutamax (Gibco, Thermo Fisher Scientific, Waltham, Massachusetts, US, #35050-038) on surfaces pre-coated with 100 μgoml^-1^ poly-L-lysine (PLL, Sigma-Aldrich, Burlington, Massachusetts, US, #P4707) and 10 μg·ml^-1^ laminin (Sigma-Aldrich, Burlington, Massachusetts, US, #L2020), unless stated otherwise. The medium was replaced four hours later by cell culture medium consisting of Neurobasal-A medium (Gibco, Thermo Fisher Scientific, Waltham, Massachusetts, US, #12348-017) modified with B27 (Gibco, Thermo Fisher Scientific, Waltham, Massachusetts, US, #17504-044), 2 mM Glutamax (Gibco, Thermo Fisher Scientific, Waltham, Massachusetts, US, #35050-038), 50 IU·ml^-1^ penicillin, 50 μg·ml^-1^ streptomycin and 2.5 μM AraC (Sigma-Aldrich, Burlington, Massachusetts, US, #C1768).

For DRGs neurons, mice pups (P3 stage) were used. After sacrifice, 20-30 DRGs were collected per animal. DRGs were then digested in a solution of DPBS supplemented with 0.03% collagenase from *Clostridium histolyticum* (Sigma-Aldrich, Burlington, Massachusetts, US, #C7657), 0.3% dispase II protease (Sigma-Aldrich, Burlington, Massachusetts, US, #D4693) and 0.18% glucose (Sigma-Aldrich, Burlington, Massachusetts, US, #G7021). The digesting DRGs were incubated with DPBS containing 0.01% deoxyribonuclease I from bovine pancreas (Sigma-Aldrich, Burlington, Massachusetts, US, #DN25) and 0.05% trypsin inhibitor from Glycine max (Sigma-Aldrich, Burlington, Massachusetts, US, #T9003). After the chemical digestion, DRGs were mechanically dissociated with a fire-polished glass Pasteur pipette. Cells were resuspended in cell growth medium modified with 100 ng·ml^-1^ NGF (Sigma-Aldrich, Burlington, Massachusetts, US, #N5415) and seeded on surfaces pre-coated with 100 μg·ml^-1^ poly-D-lysine (PDL, Sigma-Aldrich, Burlington, Massachusetts, US, #A-003-E) and 10 μg·ml^-1^ laminin, unless otherwise stated.

Half of the medium was replaced every 2-3 days. Cell cultures were maintained at 37°C in a saturated humidity atmosphere containing 95% air and 5% CO_2_.

The isolation of primary *C. elegans* neurons was performed following a previously described method (Das et al., 2021). Briefly, after synchronization, worms were seeded onto peptone-enriched plates and incubated at room temperature (RT). Once populated with eggs, the plates were washed with Milli-Q H_2_O and the eggs collected. The obtained pellet of worms and eggs was lysed through resuspension in a freshly prepared bleaching solution and rocked gently by hand for up to 10 minutes. The progress of the lysis reaction was monitored until approximately 70% of the worms were lysed. The reaction was then stopped using an egg buffer [118 mM NaCl, 48 mM KCl, 2 mM CaCl_2_, 2 mM MgCl_2_, 25 mM Hepes (pH 7.3), and an osmolarity of 340 mOsm]. After three washes with fresh egg buffer, the eggs were separated using a final 30% concentration of sucrose by centrifuging at 1200 rpm for 20 minutes. After collection and three washes with egg buffer, the eggs were treated with 0.5 U·ml^-1^ chitinase (Sigma-Aldrich, Burlington, Massachusetts, US, #C6137). To digest the eggshells, chitinase was rocked gently at RT for 40 minutes. Once approximately 80% of the eggs have been digested, the reaction was stopped with L15 medium (Sigma-Aldrich, Burlington, Massachusetts, US, #L4386 with the addition of 10% (v/v) heat-inactivated FBS, 50 U·ml^-1^ penicillin, and 50 μg·ml^-1^ streptomycin). The resulting solution passed through a 25 gauge needle 10 to 15 times. The obtained single-cell suspension was filtered through a 5 μm Durapore filter (Millipore, Burlington, Massachusetts, US, #SVLP04700) and centrifuged for 3 minutes at 3200 rpm. Cells were cultured on peanut lectin (1:10, Medicago, Denmark, #05-0116-10) coated surfaces at a density of ~ 300 cells mm^-2^ and incubated at 25 °C (unless stated otherwise). After four hours, new medium was added. Fresh medium was changed every 24 hours.

#### Nanoparticles

MNPs are magnetite nanoparticles (Fluid-MAG-ARA, Chemicell, Germany; #4115) characterized by a core of iron oxide (approximately 75±10 nm in diameter) and an outer layer of glucoronic acid. The hydrodynamic diameter is 100 nm. The saturation magnetization of 59 A·m^2^·kg^-1^, as stated from the supplier. MNPs have been added to the cell growth medium at the concentration of 5 μg·ml^-1^.

#### Microfluidic chambers

Cells were grown in microfluidic chambers assembled by mounting XONA microfluidic devices (XONA, Research Triangle Park, North Caroline, US, #RD150) on glass coverslips (22 mm in diameter). In order to promote axons to invade the axonal compartment, for hippocampal neurons, a differential coating was performed (standard coating in the somato-dendritic compartment and 500 μg·ml^-1^ PLL and 100 μg·ml^-1^ laminin for the axonal compartment) while, for DRG neurons, a NGF gradient was created (100 and 50 ng·ml^-1^ for the axonal and somato-dendritic compartment, respectively).

#### Magnetic nano-pulling assay

At DIV0, hippocampal or DRG neurons were seeded in microfluidic chambers at a density of 200,000 cells, while *C. elegans* touch receptor neurons in 35 mm glass-bottom Petri dishes #1.5 (Willcow Glass, The Netherlands, #HBST-3512) at a density of approximately 500,000 cells. Cell growth medium was modified with 5 μg·ml^-1^ MNPs four hours after seeding in hippocampal and touch receptor neurons, and the day after (DIV1) in DRGs neurons. The magnetic field (stretch group) or a null magnetic field (control group) was applied from DIV1 (if not stated differently). A Halbach-like cylinder magnetic applicator was used, which provided a constant magnetic field gradient of 46.5 T·m^-1^ in the radial centrifugal direction (Raffa et al., 2018; Riggio et al., 2014). For axonal transport studies, a neodymium disc magnet (grade N45, diameter 12 mm, height 2 mm) was used and applied for 12 hours before observations.

Hippocampal neurons were fixed in 2% PFA for 10 minutes at RT at DIV6 (if not stated differently). DRGs and touch receptor neurons were not fixed.

#### RNA extraction and quantification

The experiment was performed in microfluidic chambers and the RNA was separately extracted from the somato-dendritic and axonal compartment as described in (Nijssen et al., 2018). The nucleoSpin RNA PLUS XS Kit (Machery-Nagel, Düren, Germany, #740990.50) was used for RNA extraction. Axonal compartment was lysed without affecting the soma compartment (and vice-versa) by generating a liquid head that counteracts diffusion (Mills et al., 2018; Nijssen et al., 2018; Taylor et al., 2005). Samples were collected in lysis solution and incubated at 4°C. Lysed samples were then treated to digest the DNA, and the RNA was eluted in RNAse-free water. The Quant-iT RiboGreen RNA Kit (Thermo Fisher Scientific, Waltham, Massachusetts, US, #R11490) was used for the quantification of total RNA. QuantiTect Reverse Transcription Kit (Qiagen, Hilden, Germany, #205311) was used to produce the cDNA via reverse transcription. qPCR was performed using GoTaq®pPCR (Promega, Madison, Wisconsin, US, #A6001). The expression of two markers, the nuclear histone H1 and a ubiquitous β-actin was evaluated to rule out the presence of somas in the axonal compartment (Key Resource Table and supplementary Table S1).

#### RNA sequencing

The experiment was performed for stretch group and control group on two cellular components (axonal and somato-dendritic compartment), and in six biological replicates. In total, 24 RNA extracts were analyzed. Quality check (QC) was performed by calculating the ribosomal content (RNA integrity number). RNAseq was performed by GENOME Scan (The Netherlands) with the platform Illumina NovaSeq6000 sequencer. The RNA library was prepared using the polyA selection library and sequencing mode PE (paired end), read length 150 bp, ~ 9 Gb and 30 million XP reads per sample. Default genomes and mouse annotation were Ensembl GRCm38.p6 (available on https://www.ncbi.nlm.nih.gov/assembly/GCF_000001635.26/). Raw data quality was analysed using FastQC v0.11.9, MultiQC v1.12 QC tools (Fig. S3). Sequence reads were trimmed to remove possible adapter sequences using fastp v0.20.1 with default settings. The reads were mapped against the reference sequence using STAR v2.7.08a with default settings (Fig. S4) and the number of reads per gene was determined with HTSeq v0.13.5 and the Ensembl GRCm38.p6 GTF annotation file (Fig. S5). Differential gene expression analysis was performed using DESeq2 v 1.30.1. Datasets are available in the public repository GEO (accession number: GSE197808).

#### Quality check at the biological and bioinformatic level

In the microfluidic chamber model, only axons should be able to spread into the axonal compartment, due to the small cross-section of the microgrooves. However, to ensure the absence of any somatic cross-contamination, samples were inspected for the presence of nuclei in the axonal compartment, all of which were negative to Hoechst staining (n=6) (Figure S1). To further rule out additional sources of contamination (*e.g*. spillage of lysis buffer between the two compartments), a qPCR analysis was performed on RNA extracts for the detection of a nuclear marker, the histone protein H1. β-actin mRNA (housekeeping) was detected in both compartments while H1 mRNA was present in the somato-dendritic compartment but not in the axonal one (n=6), which meant that the cross-contamination was negligible (Table S1). As a further quality control, we decided to eliminate axon samples with trace levels of soma contamination at the bioinformatic level. Specifically, we performed a PCA based on all expressed genes. PC1 reflected the major variance (85% and 3% of variance associated with PC1 and PC2, respectively, Fig. 6A). PC1 and the sample-to-sample distance map (not shown) confirmed that samples extracted from the axonal compartment cluster and segregate from samples extracted from the somato-dendritic one based on the gene expression patterns. One axon sample, out of 12, was intermediate (marked with a circle, Fig. 6A). To confirm the presence of a cross-contamination for this sample, the number of detected genes was counted. Excluding this sample from the count of detected genes, axonal samples contained on average 6,382±2025 detected genes (n=11), and somato-dendritic samples contained 23,329±753 detected genes in total (n=12) (*t* test, two-tailed, *p*<0.0001). The sample marked with a circle in Fig. 6A contained 14159 detected genes, which is clearly an intermediate value between somato-dendritic and axonal samples, suggesting a contamination by soma. This sample, probably contaminated by one or more somas, was then discarded from the subsequent analysis.

#### GO and GOEA

GO analysis was carried out using DAVID version 6.8 (https://david.ncifcrf.gov/) and the subdatabases GOTERM_CC_DIRECT and the feature “functional annotation chart” (Huang et al., 2009). GOEA was conducted using SynGO (https://www.syngoportal.org/) to perform enrichment analysis of synaptic genes in the function domain (Koopmans et al., 2019)).

#### Ribopuromycylation

The study of ribosomes in active translation was performed through the ribopuromycylation (RPM method, modified from (Bastide et al., 2018)). Briefly, hippocampal neurons were cultured in microfluidic chambers from DIV0 and stimulated from DIV1 to DIV6. At DIV6, neurons were treated with 200 μM emetine (Sigma-Aldrich, Burlington, Massachusetts, US, #E2375) and 100 μM puromycin (Sigma-Aldrich, Burlington, Massachusetts, US, #P7255) for 10 minutes at 37°C. Samples were then washed with ice-cold 0.0003% digitonin (Sigma-Aldrich, Burlington, Massachusetts, US; #D141) for two minutes. Lastly, samples were washed with ice-cold DPBS and fixed in 2% PFA, 7.5% sucrose (Sigma-Aldrich, Burlington, Massachusetts, US, #S0389) for 20 minutes at RT.

#### Immunostaining and imaging

For all the experiments (except those related to Fig. 1C4, 1D4 and Fig. 3), after fixation, hippocampal neurons were permeabilized in 0.5% Triton X-100 for 10 minutes at RT. Samples were then blocked in 5% serum, 0.3% Triton X-100 in DPBS for 1 hour at RT. Primary antibodies were diluted in 3% serum, 0.2% Triton X-100 in DPBS as follows: TUBB3 (Sigma-Aldrich, Burlington, Massachusetts, US; #T8578, 1:500) Synapsin I (Synaptic Systems, Goettingen, Germany, #106 103, 1:500), Homer 1b/c (Synaptic Systems, Goettingen, Germany, #106 023, 1:350), VGlut1 (Sigma-Aldrich, Burlington, Massachusetts, US; # AMAB91041, 1:500). After overnight incubation at 4°C, samples were washed and then incubated with secondary antibodies (Thermo Fisher Scientific, Waltham, Massachusetts, US, #O6380, #A32731, #A11011, #A32728, #R6393, #A21449, 1:500; Abcam, Cambridge, UK, #ab150169, 1:500) and Hoechst 33342 (Thermo Fisher Scientific, Waltham, Massachusetts, US, #H3570, 1:1000) for 1 hour at RT. For experiments related to Fig. 1C4, Fig. 1D4 and Fig. 3, we followed a protocol modified from Cioni and colleagues (Cioni et al., 2019). Briefly, at DIV6, samples were fixed in 2% PFA, 7.5% sucrose in DPBS for 20 min at RT. Samples were then washed with DPBS and Triton X-100 in very low concentration (0.001% in DPBS). Samples were permeabilized with a solution of 0.1% Triton X-100 in DPBS for 5 minutes, and blocked with 5% goat serum in DPBS for 30 minutes. Samples were incubated with primary antibodies (TUBB3, Sigma-Aldrich, Burlington, Massachusetts, US, #T8578, 1:500; TUBB3, Abcam, Cambridge, UK, #ab41489, 1:1000; Mark4, Abcam, Cambridge, UK, # ab124267, 1:200; tau ser262, Abcam, Cambridge, UK, #ab131354, 1:200; S6, Cell Signaling, Danvers, Massachusetts, US, #2217, 1:200; Rab7, Abcam, Cambridge, UK, #ab137029, 1:200; anti-Puromycin, Sigma-Aldrich, Burlington, Massachusetts, US, #MABE343, 1:1000) at 4°C overnight. Then samples were incubated with secondary antibodies (see above) and Syto RNA select (Thermo Fisher Scientific, Waltham, Massachusetts, US, #S32703, 1:2500), or Hoechst 33342 (see above) for 45 minutes at RT. Samples were imaged with a laser scanning confocal microscope (Nikon A1, Eclipse Ti). Images were acquired with a 60x objective oil immersion. Series of □ 30 optical plans in Z were acquired at 1024 × 1024 pixel resolution with a z-step of 0.2 μm. Images were acquired using a 405 nm laser (425-475 emission filter) or a 488 nm laser (500-550 emission filter) or a 561 laser (570-620 emission filter) or a 640 nm laser (663-738 emission filter).

Before imaging the *C. elegans* neurons, the medium was changed. Image acquisition was performed using an inverted microscope (DMi8, Leica), either with a 40×/1.1 water immersion lens or a 25×/0.95 water immersion lens. A multi-wavelength LED light source (SpectraX, Lumencor) was used for fluorescence excitation. Touch receptor neurons expressing calcium-sensitive GCaMP6s and tag-RFP were excited with 470/24 nm and 550/15 nm band-pass filtered LEDs at 8-15% and 20-30% of the output power, respectively (measured 4-7 mW and 14-21 mW at the sample plane with a microscope slide power meter, Thorlabs S170C). Excitation power was kept constant for both the control and stretched conditions of the same replicate. Fluorescence emission was directed to an sCMOS camera (Hamamatsu Orca Flash 4 V3) with a quad-edge dichroic splitter (Semrock, FF409/493/573/652-Di02-25×36). Green and red fluorophore signals were later separated with an image splitting unit (Hamamatsu, W-view Gemini A12801-01), in which a 538 nm edge dichroic splitter (Semrock, FF528-FDi1-25-36), and 512/25 nm (Semrock, FF01-512/25-25) and 670/30 nm (FF01-670/30-25) band-pass emission filters were built. The sCMOS camera was used in W-view mode, enabling simultaneous dual fluorophore imaging at an exposure time of 200 ms for both top (GCaMP) and bottom (tag-RFP) camera halves. Motor neurons expressing solely GCaMP6s were directly imaged with no image splitting unit.

#### Sample preparation for TEM

For the ultrastructural characterization, hippocampal neurons were treated as previously described (Convertino et al., 2020). Briefly, neurons were fixed with an aldehydic solution (1.5% glutaraldehyde in 0.1 M Cacodylate buffer - pH 7.4), washed, and post-fixed with reduced osmium tetroxide solution (1% K_3_Fe(CN)_6_ + 1% OsO_4_ in 0.1 M Cacodylate buffer). After rinses, neurons were stained with our homemade staining solution (X solution diluted 1:10 (v/v) in 20% ethanol/water (Moscardini et al., 2020), then dehydrated, with an increasing series of ethanol concentrations. Samples were then embedded in epoxy resin (Epoxy embedding medium kit, Merck KGaA, Darmstadt, Germany) which was then baked for 48 hours at 60°C. After the coverslips have been removed, embedded neurons were mounted on a resin support and sectioned with a UC7 ultramicrotome (Leica Microsystems, Vienna, Austria) with a 35° diamond knife (Diatome Ltd, Switzerlands). Sections of 80 nm were collected on 300 mesh copper grids (G300Cu - Electron Microscopy Science, Hatfield, PA, USA).

Grids were analyzed with a Zeiss Libra 120 Plus transmission electron microscope, operating at 120 kV, and equipped with an in-column omega filter (for the energy filtered imaging).

#### NGF single vesicle tracking

fluoNGF was obtained by performing an enzymatic fluorolabeling reaction on the YBBR-tagged NGF recombinantly produced in E. coli (di Matteo et al., 2017). A total of 90-100 mg purified YBBR-NGF (kindly donated by Prof. A. Cattaneo, Bio@SNS Lab, Pisa) was incubated with 73 μM CoA-Alexa647, 17μM Sfp Synthase, 36 mM MgCl_2_ in DPBS up to 270 μl final volume for 30 min at 37°C; the excess of free fluorophore was removed by desalt spin-column and the fluoNGF obtained was stored at 4°C for a maximum of 10 days. This reaction enabled the NGF C-terminus to be stoichiometrically labelled with Alexa647 fluorophore using a method first described in Yin and colleagues (Yin et al., 2005) and optimized as in (Amodeo et al., 2020), to finally achieve the covalent binding of two fluorophores per neurotrophin dimer. At DIV3 (after overnight stimulation), DRG cultures were prepared for time-lapse studies. To track the vesicles, 2 nM fluoNGF was applied in the axon side, incubated for 1 hour at 37°C, after which the axonal medium was replaced.

In order to acquire time-lapse videos, we used an inverted epi-fluorescence microscope (Leica AF6000) equipped with Leica TIRF-AM module, incubator chamber at 37°C, 5% CO_2_, an electron multiplying charge-coupled device camera (ImagEM C9100-13, Hamamatsu), and a 100× oil immersion objective (NA 1.47), which enabled the acquisition of fields of 512×512pixels (116.80×116.80 μm) typically comprising two microchannels. fluoNGF vesicles were imaged inside the microchannels in epifluorescence configuration, using a 635 nm laser line at maximum power, a Cy5 Leica1152303 fluorescence cube and an exposure time of 100 ms. Up to 1000 frames for each time-lapse video were acquired, and each chip was imaged for about 45 min.

### Quantification and statistical analysis

#### Image analysis

NeuronJ, a Fiji plugin, was used to evaluate axon length (Meijering et al., 2004). Briefly, axons (TUBB3 stained) were traced with the tracing tool by choosing the exit point from the microchannels as the starting point. For *C. elegans* neurons, the green GCaMP channel was used to determine the length of the axon.

For NGF vesicle tracking analysis, scripts in MatLab (The MathWorks) were used in order to detect single vesicles containing fluoNGF along the axon and to track them, so as to assign them a dynamic mode based on the direction of movement and velocity, as described in (Convertino et al., 2020). Kymograph analysis was carried out using ImageJ image analysis software.

For fluorescence quantification, we evaluated the total fluorescence (*f*) and/or the mean fluorescence 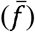, defined as:

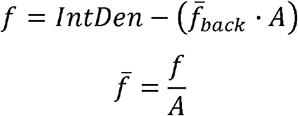

where *IntDen* is the integrated density (defined as the sum of all the pixel intensities in that selected region), A is the area of the ROI (region of interest) and 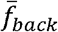 is the mean fluorescence of background readings.

Co-localization studies between late endosomes and RNA granules or active ribosomes were performed with Diana, a Fiji plugin (Gilles et al., 2017). Taking advantage of the integrated features of the plugin, each channel was first segmented (default parameters for LE and active ribosomes; min. objects size = 1 pixel for LE and RNA granules), then the percentage of co-localization between the two objects of interest (LE and RNA granules or LE and active ribosomes) was evaluated.

The Diana plugin was also exploited for the evaluation of RNA granules density. Briefly, images were segmented in Diana (min. objects size = 1 pixel) and the total volume of the RNA granules contained in one neurite was collected and then normalized for the corresponding area. The same plugin was also used to study the number, volume and fluorescence (as integrated density) of synapsin I spots contained in a defined neurite compartment. After the selection of the region of interest, the synapsin I channel was segmented with default parameters. The values related to the mentioned parameters were then collected and normalized for the analyzed area.

To estimate the co-localization between the spots of the pre-synaptic marker VGlut1 and the post-synaptic marker homer 1b/c, the plugin puncta analyzer was applied, as previously described (Ippolito and Eroglu, 2010). Briefly, both channels were converted using the maximum intensity of the Z projection and merged. Following the identification of a specific ROI corresponding to all the neurites of the cell, and keeping default parameters, the plugin quantifies the puncta in each channel and the co-localized puncta between the two channels. The value obtained was then normalized for the considered area.

#### TEM analysis

TEM analysis was carried out using ImageJ and the plugin NeuronJ (Meijering et al., 2004). For MT density, the MTs, recognized as tubular structures, were counted manually in a selected and organelle-free region of the neurite (Fig. S6, dashed yellow lines). The diameter of the corresponding region was then measured, and the number of MTs normalized, obtaining one value per neurite. To study the endoplasmic reticulum (ER), the cisternae were traced with NeuronJ and then normalized for the considered neurite area (Fig. S6, magenta lines). NeuronJ was also exploited for the analysis of the post-synaptic density (PSD) region. Specifically, it was used to measure the length of PSDs (Fig. S6, white line) in DIV14 control and stretched samples, and the mean value was considered for each synapse. The number of PSDs of each synapse was also counted manually in the two conditions. To determine the mitochondria density, mitochondria were counted manually (Fig. S6, white “*”) in DIV14 stretched and unstretched samples. The corresponding number was then normalized for the total area of the neurite, obtaining one value per neurite.

#### Statistical analysis

Data were plotted with GraphPad 7.0. Values are reported as the mean ± standard error of the mean (SEM). The normality of the distribution was tested using the D’Agostino & Pearson normality test, Shapiro-Wilk normality test, or Kolmogorov-Smirnov normality test. For normally distributed data, we used the *t* test for unpaired data followed by Bonferroni correction or two-way ANOVA test. For non-normally distributed data, the Mann-Whitney test or Kruskal-Wallis test with post hoc Dunn’s test, were carried out. Significance was set at *p* ≤ 0.05.

### Data and code availability

All data are available in the manuscript or supplemental information. The codes used have been previously published and are freely available (see Key Resources Table). Any additional information required to reanalyze the data reported in this paper is available from the lead contact upon request.

### Supplemental information

A primary supplemental file that contains all supplemental figures and table is provided.

Multimedia files:

- Movie S1: imaging and tracking of NGF vesicles in DRG control axons;
- Movies S2: imaging and tracking of NGF vesicles in DRG stretched axons;
- Movies S3: imaging and tracking of NGF vesicles in DRG stretched axons, 30 minutes after the magnet had been removed.

## Notes

https://drive.google.com/drive/folders/17BE4l_ZO2PMRB3FEUbkX_2qti8Zd_HgR?usp=sharing

### Competing Interest Statement

The authors have declared no competing interest.

## References

Ahmed, W.W., and Saif, T.A. (2015). Active transport of vesicles in neurons is modulated by mechanical tension. Scientific Reports 4, 4481. https://doi.org/10.1038/srep04481.

Akiyoshi, B., Sarangapani, K.K., Powers, A.F., Nelson, C.R., Reichow, S.L., Arellano-Santoyo, H., Gonen, T., Ranish, J.A., Asbury, C.L., and Biggins, S. (2010). Tension directly stabilizes reconstituted kinetochore-microtubule attachments. Nature 468, 576–579. .

Amodeo, R., Convertino, D., Calvello, M., Ceccarelli, L., Bonsignore, F., Ravelli, C., Cattaneo, A., Martini, C., Luin, S., Mitola, S., et al. (2020). Fluorolabeling of the PPTase-Related Chemical Tags: Comparative Study of Different Membrane Receptors and Different Fluorophores in the Labeling Reactions. Frontiers in Molecular Biosciences 7.https://doi.org/10.3389/fmolb.2020.00195.

Bastide, A., Yewdell, J., and David, A. (2018). The RiboPuromycylation Method (RPM): an Immunofluorescence Technique to Map Translation Sites at the Sub-cellular Level. BIO-PROTOCOL 8. https://doi.org/10.21769/BioProtoc.2669.

Berger, F., Klumpp, S., and Lipowsky, R. (2019). Force-Dependent Unbinding Rate of Molecular Motors from Stationary Optical Trap Data. Nano Letters 19, 2598–2602. https://doi.org/10.1021/acs.nanolett.9b00417.

Biernat, J., Gustke, N., Drewes, G., and Mandelkow, E. (1993). Phosphorylation of Ser262 strongly reduces binding of tau to microtubules: distinction between PHF-like immunoreactivity and microtubule binding. Neuron 11, 153–163. .

Bray, D. (1984). Axonal growth in response to experimentally applied mechanical tension. Dev Biol 102, 379–389. .

Chowdary, P.C., McGuire, A., Lee, Y., Che, D., Hanson, L., Osakada, Y., Ooi, C., Xie, C., Wang, S.X., and Cui, B. (2019). Magnetic manipulation of axonal endosome transport in live neurons. BioRxiv 733253. .

Chowdary, P.D., Xie, C., Osakada, Y., Che, D.L., and Cui, B. (2013). Magnetic Manipulation of Axonal Transport in Live Neurons. Biophysical Journal 104, 652a. https://doi.org/10.1016/j.bpj.2012.11.3600.

Cioni, J.-M., Lin, J.Q., Holtermann, A. v., Koppers, M., Jakobs, M.A.H., Azizi, A., Turner-Bridger, B., Shigeoka, T., Franze, K., Harris, W.A., et al. (2019). Late Endosomes Act as mRNA Translation Platforms and Sustain Mitochondria in Axons. Cell 176, 56–72.e15. https://doi.org/10.1016/j.cell.2018.11.030.

Convertino, D., Fabbri, F., Mishra, N., Mainardi, M., Cappello, V., Testa, G., Capsoni, S., Albertazzi, L., Luin, S., Marchetti, L., et al. (2020). Graphene Promotes Axon Elongation through Local Stall of Nerve Growth Factor Signaling Endosomes. Nano Letters 20, 3633–3641. https://doi.org/10.1021/acs.nanolett.0c00571.

Dai, R., Hang, Y., Liu, Q., Zhang, S., Wang, L., Pan, Y., and Chen, H. (2019). Improved neural differentiation of stem cells mediated by magnetic nanoparticle-based biophysical stimulation. Journal of Materials Chemistry B 7, 4161–4168. https://doi.org/10.1039/C9TB00678H.

Falconieri, A., Taparia, N., de Vincentiis, S., Cappello, V., Sniadecki, N.J., and Raffa, V. (2021). MAGNETICALLY-ACTUATED MICROPOSTS STIMULATE AXON GROWTH. Biophysical Journal https://doi.org/10.1016/j.bpj.2021.12.041.

Farías, G.G., Fréal, A., Tortosa, E., Stucchi, R., Pan, X., Portegies, S., Will, L., Altelaar, M., and Hoogenraad, C.C. (2019). Feedback-Driven Mechanisms between Microtubules and the Endoplasmic Reticulum Instruct Neuronal Polarity. Neuron 102, 184–201.e8. https://doi.org/10.1016/j.neuron.2019.01.030.

Franck, A.D., Powers, A.F., Gestaut, D.R., Gonen, T., Davis, T.N., and Asbury, C.L. (2007). Tension applied through the Dam1 complex promotes microtubule elongation providing a direct mechanism for length control in mitosis. Nat Cell Biol 9, 832–837. .

Franze, K. (2013). The mechanical control of nervous system development. Development 140, 3069–3077. .

Fukushige, T., Siddiqui, Z.K., Chou, M., Culotti, J.G., Gogonea, C.B., Siddiqui, S.S., and Hamelin, M. (1999). MEC-12, an alpha-tubulin required for touch sensitivity in C. elegans. Journal of Cell Science 112, 395–403. https://doi.org/10.1242/jcs.112.3.395.

Gahl, T.J., and Kunze, A. (2018). Force-mediating magnetic nanoparticles to engineer neuronal cell function. Frontiers in Neuroscience https://doi.org/10.3389/fnins.2018.00299.

Gangatharan, G., Schneider-Maunoury, S., and Breau, M.A. (2018). Role of mechanical cues in shaping neuronal morphology and connectivity. Biology of the Cell 110, 125–136. https://doi.org/10.1111/boc.201800003.

Gennerich, A., Carter, A.P., Reck-Peterson, S.L., and Vale, R.D. (2007). Force-Induced Bidirectional Stepping of Cytoplasmic Dynein. Cell 131, 952–965. https://doi.org/10.1016/j.cell.2007.10.016.

Gilles, J.-F., dos Santos, M., Boudier, T., Bolte, S., and Heck, N. (2017). DiAna, an ImageJ tool for object-based 3D co-localization and distance analysis. Methods 115, 55–64. https://doi.org/10.1016/j.ymeth.2016.11.016.

Gitler, D., Takagishi, Y., Feng, J., Ren, Y., Rodriguiz, R.M., Wetsel, W.C., Greengard, P., and Augustine, G.J. (2004). Different presynaptic roles of synapsins at excitatory and inhibitory synapses. Journal of Neuroscience 24, 11368–11380. .

Goedert, M., Baur, C.P., Ahringer, J., Jakes, R., Hasegawa, M., Spillantini, M.G., Smith, M.J., and Hill, F. (1996). PTL-1, a microtubule-associated protein with tau-like repeats from the nematode Caenorhabditis elegans. Journal of Cell Science 109, 2661–2672. https://doi.org/10.1242/jcs.109.11.2661.

Guedes-Dias, P., and Holzbaur, E.L.F. (2019). Axonal transport: Driving synaptic function. Science (1979) 366. .

Gürth, C.-M., Dankovich, T.M., Rizzoli, S.O., and D’Este, E. (2020). Synaptic activity and strength are reflected by changes in the post-synaptic secretory pathway. Scientific Reports 10, 20576. https://doi.org/10.1038/s41598-020-77260-2.

Hamant, O., Inoue, D., Bouchez, D., Dumais, J., and Mjolsness, E. (2019). Are microtubules tension sensors? Nature Communications 10, 2360. https://doi.org/10.1038/s41467-019-10207-y.

Holt, C.E., Martin, K.C., and Schuman, E.M. (2019). Local translation in neurons: visualization and function. Nature Structural & Molecular Biology 26, 557–566. https://doi.org/10.1038/s41594-019-0263-5.

Howe, C.L., and Mobley, W.C. (2004). Signaling endosome hypothesis: A cellular mechanism for long distance communication. Journal of Neurobiology 58, 207–216. https://doi.org/10.1002/neu.10323.

Huang, D.W., Sherman, B.T., and Lempicki, R.A. (2009). Systematic and integrative analysis of large gene lists using DAVID bioinformatics resources. Nature Protocols 4, 44–57. https://doi.org/10.1038/nprot.2008.211.

Inoue, D., Nitta, T., Kabir, A.M.R., Sada, K., Gong, J.P., Konagaya, A., and Kakugo, A. (2016). Sensing surface mechanical deformation using active probes driven by motor proteins. Nat Commun 7, 1–10. .

Ippolito, D.M., and Eroglu, C. (2010). Quantifying Synapses: an Immunocytochemistry-based Assay to Quantify Synapse Number. Journal of Visualized Experiments https://doi.org/10.3791/2270.

Kadavath, H., Hofele, R. V, Biernat, J., Kumar, S., Tepper, K., Urlaub, H., Mandelkow, E., and Zweckstetter, M. (2015). Tau stabilizes microtubules by binding at the interface between tubulin heterodimers. Proceedings of the National Academy of Sciences 112, 7501–7506. .

Kim, H.J., Park, J.W., Park, J.W., Byun, J.H., Vahidi, B., Rhee, S.W., and Jeon, N.L. (2012). Integrated microfluidics platforms for investigating injury and regeneration of CNS axons. Ann Biomed Eng 40, 1268–1276. .

Kolomeisky, A.B., and Fisher, M.E. (2001). Force-Velocity Relation for Growing Microtubules. Biophysical Journal 80, 149–154. https://doi.org/10.1016/S0006-3495(01)76002-X.

Koopmans, F., van Nierop, P., Andres-Alonso, M., Byrnes, A., Cijsouw, T., Coba, M.P., Cornelisse, L.N., Farrell, R.J., Goldschmidt, H.L., Howrigan, D.P., et al. (2019). SynGO: An Evidence-Based, Expert-Curated Knowledge Base for the Synapse. Neuron 103, 217–234.e4. https://doi.org/10.1016/j.neuron.2019.05.002.

Krieg, M., Dunn, A.R., and Goodman, M.B. (2014). Mechanical control of the sense of touch by β-spectrin. Nature Cell Biology 16, 224–233. https://doi.org/10.1038/ncb2915.

Krieg, M., Stühmer, J., Cueva, J.G., Fetter, R., Spilker, K., Cremers, D., Shen, K., Dunn, A.R., and Goodman, M.B. (2017). Genetic defects in β-spectrin and tau sensitize C. elegans axons to movement-induced damage via torque-tension coupling. Elife 6. https://doi.org/10.7554/eLife.20172.

Kunze, A., Tseng, P., Godzich, C., Murray, C., Caputo, A., Schweizer, F.E., and di Carlo, D. (2015). Engineering cortical neuron polarity with nanomagnets on a chip. ACS Nano https://doi.org/10.1021/nn505330w.

Kunze, A., Murray, C.T., Godzich, C., Lin, J., Owsley, K., Tay, A., and Di Carlo, D. (2017). Modulating motility of intracellular vesicles in cortical neurons with nanomagnetic forces on-chip. Lab on a Chip 17, 842–854. .

Lamoureux, P., Buxbaum, R.E., and Heidemann, S.R. (1989). Direct evidence that growth cones pull. Nature 340, 159–162. .

Lamoureux, P., Heidemann, S.R., Martzke, N.R., and Miller, K.E. (2010). Growth and elongation within and along the axon. Dev Neurobiol 70, 135–149. .

Lee, A.C., and Suter, D.M. (2008). Quantitative analysis of microtubule dynamics during adhesion mediated growth cone guidance. Dev Neurobiol 68, 1363–1377. .

Leidel, C., Longoria, R.A., Gutierrez, F.M., and Shubeita, G.T. (2012). Measuring Molecular Motor Forces In Vivo: Implications for Tug-of-War Models of Bidirectional Transport. Biophysical Journal 103, 492–500. https://doi.org/10.1016/j.bpj.2012.06.038.

Li, G., and Moore, J.K. (2020). Microtubule dynamics at low temperature: evidence that tubulin recycling limits assembly. Molecular Biology of the Cell 31, 1154–1166. https://doi.org/10.1091/mbc.E19-11-0634.

Liao, Y.-C., Fernandopulle, M.S., Wang, G., Choi, H., Hao, L., Drerup, C.M., Patel, R., Qamar, S., Nixon-Abell, J., Shen, Y., et al. (2019). RNA Granules Hitchhike on Lysosomes for Long-Distance Transport, Using Annexin A11 as a Molecular Tether. Cell 179, 147–164.e20. https://doi.org/10.1016/j.cell.2019.08.050.

Lin, M.-Y., and Sheng, Z.-H. (2015). Regulation of mitochondrial transport in neurons. Experimental Cell Research 334, 35–44. https://doi.org/10.1016/j.yexcr.2015.01.004.

Lindwall, G., and Cole, R.D. (1984). Phosphorylation affects the ability of tau protein to promote microtubule assembly. Journal of Biological Chemistry 259, 5301–5305. .

Liu, F., Li, B., Tung, E., Grundke Iqbal, I., Iqbal, K., and Gong, C. (2007). Site specific effects of tau phosphorylation on its microtubule assembly activity and self aggregation. European Journal of Neuroscience 26, 3429–3436. .

Liu, Y.-T., Tao, C.-L., Lau, P.-M., Zhou, Z.H., and Bi, G.-Q. (2019). Postsynaptic protein organization revealed by electron microscopy. Current Opinion in Structural Biology 54, 152–160. https://doi.org/10.1016/j.sbi.2019.02.012.

Lockhead, D., Schwarz, E.M., O’Hagan, R., Bellotti, S., Krieg, M., Barr, M.M., Dunn, A.R., Sternberg, P.W., and Goodman, M.B. (2016). The tubulin repertoire of *Caenorhabditis elegans* sensory neurons and its context-dependent role in process outgrowth. Molecular Biology of the Cell 27, 3717–3728. https://doi.org/10.1091/mbc.e16-06-0473.

Lowery, L.A., and Van Vactor, D. (2009). The trip of the tip: understanding the growth cone machinery. Nature Reviews Molecular Cell Biology 10, 332–343. .

Maeder, C.I., Shen, K., and Hoogenraad, C.C. (2014). Axon and dendritic trafficking. Curr Opin Neurobiol 27, 165–170. .

Magdesian, M.H., Lopez-Ayon, G.M., Mori, M., Boudreau, D., Goulet-Hanssens, A., Sanz, R., Miyahara, Y., Barrett, C.J., Fournier, A.E., and De Koninck, Y. (2016). Rapid mechanically controlled rewiring of neuronal circuits. Journal of Neuroscience 36, 979–987. .

Marsh, L., and Letourneau, P.C. (1984). Growth of neurites without filopodial or lamellipodial activity in the presence of cytochalasin B. J Cell Biol 99, 2041–2047. .

di Matteo, P., Calvello, M., Luin, S., Marchetti, L., and Cattaneo, A. (2017). An Optimized Procedure for the Site-Directed Labeling of NGF and proNGF for Imaging Purposes. Frontiers in Molecular Biosciences 4. https://doi.org/10.3389/fmolb.2017.00004.

Meijering, E., Jacob, M., Sarria, J.-C.F., Steiner, P., Hirling, H., and Unser, M. (2004). Design and validation of a tool for neurite tracing and analysis in fluorescence microscopy images. Cytometry 58A, 167–176. https://doi.org/10.1002/cyto.a.20022.

Merianda, T.T., Lin, A.C., Lam, J.S.Y., Vuppalanchi, D., Willis, D.E., Karin, N., Holt, C.E., and Twiss, J.L. (2009). A functional equivalent of endoplasmic reticulum and Golgi in axons for secretion of locally synthesized proteins. Molecular and Cellular Neuroscience 40, 128–142. https://doi.org/10.1016/j.mcn.2008.09.008.

Miller, K.E., and Sheetz, M.P. (2006). Direct evidence for coherent low velocity axonal transport of mitochondria. J Cell Biol 173, 373–381. .

Miller, K.E., and Suter, D.M. (2018). An Integrated Cytoskeletal Model of Neurite Outgrowth. Frontiers in Cellular Neuroscience 12. https://doi.org/10.3389/fncel.2018.00447.

Mills, R., Taylor-Weiner, H., Correia, J.C., Agudelo, L.Z., Allodi, I., Kolonelou, C., Martinez-Redondo, V., Ferreira, D.M.S., Nichterwitz, S., Comley, L.H., et al. (2018). Neurturin is a PGC-1α1-controlled myokine that promotes motor neuron recruitment and neuromuscular junction formation. Molecular Metabolism 7, 12–22. https://doi.org/10.1016/j.molmet.2017.11.001.

Moscardini, A., di Pietro, S., Signore, G., Parlanti, P., Santi, M., Gemmi, M., and Cappello, V. (2020). Uranium-free X solution: a new generation contrast agent for biological samples ultrastructure. Scientific Reports 10, 11540. https://doi.org/10.1038/s41598-020-68405-4.

Muller, D., Buchs, P.-A., and Stoppini, L. (1993). Time course of synaptic development in hippocampal organotypic cultures. Developmental Brain Research 71, 93–100. https://doi.org/10.1016/0165-3806(93)90109-N.

Nagano, S., and Araki, T. (2021). Axonal Transport and Local Translation of mRNA in Neurodegenerative Diseases. Frontiers in Molecular Neuroscience 14. https://doi.org/10.3389/fnmol.2021.697973.

Nagano, S., Jinno, J., Abdelhamid, R.F., Jin, Y., Shibata, M., Watanabe, S., Hirokawa, S., Nishizawa, M., Sakimura, K., Onodera, O., et al. (2020). TDP-43 transports ribosomal protein mRNA to regulate axonal local translation in neuronal axons. Acta Neuropathologica 140, 695–713. https://doi.org/10.1007/s00401-020-02205-y.

Nijssen, J., Aguila, J., Hoogstraaten, R., Kee, N., and Hedlund, E. (2018). Axon-Seq Decodes the Motor Axon Transcriptome and Its Modulation in Response to ALS. Stem Cell Reports 11, 1565–1578. https://doi.org/10.1016/j.stemcr.2018.11.005.

Oba, T., Saito, T., Asada, A., Shimizu, S., Iijima, K.M., and Ando, K. (2020). Microtubule affinity—regulating kinase 4 with an Alzheimer’s disease-related mutation promotes tau accumulation and exacerbates neurodegeneration. Journal of Biological Chemistry 295, 17138–17147. .

O’Toole, M., Lamoureux, P., and Miller, K.E. (2015). Measurement of Subcellular Force Generation in Neurons. Biophysical Journal 108, 1027–1037. https://doi.org/10.1016/j.bpj.2015.01.021.

Pfister, B.J., Iwata, A., Meaney, D.F., and Smith, D.H. (2004). Extreme stretch growth of integrated axons. Journal of Neuroscience 24, 7978–7983. .

Putnam, A.J., Schultz, K., and Mooney, D.J. (2001). Control of microtubule assembly by extracellular matrix and externally applied strain. American Journal of Physiology-Cell Physiology 280, C556–C564. .

Raffa, V., Falcone, F., de Vincentiis, S., Falconieri, A., Calatayud, M.P., Goya, G.F., and Cuschieri, A. (2018). Piconewton Mechanical Forces Promote Neurite Growth. Biophysical Journal 115, 2026–2033. https://doi.org/10.1016/j.bpj.2018.10.009.

Riggio, C., Calatayud, M.P., Giannaccini, M., Sanz, B., Torres, T.E., Fernández-Pacheco, R., Ripoli, A., Ibarra, M.R., Dente, L., Cuschieri, A., et al. (2014). The orientation of the neuronal growth process can be directed via magnetic nanoparticles under an applied magnetic field. Nanomedicine: Nanotechnology, Biology, and Medicine 10. https://doi.org/10.1016/j.nano.2013.12.008.

de Rooij, R., Kuhl, E., and Miller, K.E. (2018). Modeling the axon as an active partner with the growth cone in axonal elongation. Biophys J 115, 1783–1795. .

Sánchez-Huertas, C., and Herrera, E. (2021). With the Permission of Microtubules: An Updated Overview on Microtubule Function During Axon Pathfinding. Frontiers in Molecular Neuroscience 14. https://doi.org/10.3389/fnmol.2021.759404.

Scott, C.W., Spreen, R.C., Herman, J.L., Chow, F.P., Davison, M.D., Young, J., and Caputo, C.B. (1993). Phosphorylation of recombinant tau by cAMP-dependent protein kinase. Identification of phosphorylation sites and effect on microtubule assembly. Journal of Biological Chemistry 268, 1166–1173. .

Sengupta, A., Kabat, J., Novak, M., Wu, Q., Grundke-Iqbal, I., and Iqbal, K. (1998). Phosphorylation of tau at both Thr 231 and Ser 262 is required for maximal inhibition of its binding to microtubules. Arch Biochem Biophys 357, 299–309. .

Shigeoka, T., Koppers, M., Wong, H.H.-W., Lin, J.Q., Cagnetta, R., Dwivedy, A., de Freitas Nascimento, J., van Tartwijk, F.W., Ströhl, F., Cioni, J.-M., et al. (2019). On-Site Ribosome Remodeling by Locally Synthesized Ribosomal Proteins in Axons. Cell Reports 29, 3605–3619.e10. https://doi.org/10.1016/j.celrep.2019.11.025.

Spillane, M., Ketschek, A., Merianda, T.T., Twiss, J.L., and Gallo, G. (2013). Mitochondria Coordinate Sites of Axon Branching through Localized Intra-axonal Protein Synthesis. Cell Reports 5, 1564–1575. https://doi.org/10.1016/j.celrep.2013.11.022.

Steketee, M.B., Moysidis, S.N., Jin, X.L., Weinstein, J.E., Pita-Thomas, W., Raju, H.B., Iqbal, S., and Goldberg, J.L. (2011). Nanoparticle-mediated signaling endosome localization regulates growth cone motility and neurite growth. Proc Natl Acad Sci U S A https://doi.org/10.1073/pnas.1019624108.

Steketee, M.B., Oboudiyat, C., Daneman, R., Trakhtenberg, E., Lamoureux, P., Weinstein, J.E., Heidemann, S., Barres, B.A., and Goldberg, J.L. (2014). Regulation of Intrinsic Axon Growth Ability at Retinal Ganglion Cell Growth Cones. Investigative Opthalmology & Visual Science 55, 4369. https://doi.org/10.1167/iovs.14-13882.

Suter, D.M., and Miller, K.E. (2011). The emerging role of forces in axonal elongation. Progress in Neurobiology 94, 91–101. https://doi.org/10.1016/j.pneurobio.2011.04.002.

Tamariz, E., and Varela-Echavarría, A. (2015). The discovery of the growth cone and its influence on the study of axon guidance. Front Neuroanat 9, 51. .

Tay, A., Kunze, A., Murray, C., and di Carlo, D. (2016). Induction of Calcium Influx in Cortical Neural Networks by Nanomagnetic Forces. ACS Nano 10, 2331–2341. https://doi.org/10.1021/acsnano.5b07118.

Taylor, A.M., Blurton-Jones, M., Rhee, S.W., Cribbs, D.H., Cotman, C.W., and Jeon, N.L. (2005). A microfluidic culture platform for CNS axonal injury, regeneration and transport. Nature Methods 2, 599–605. https://doi.org/10.1038/nmeth777.

Toriyama, M., Kozawa, S., Sakumura, Y., and Inagaki, N. (2013). Conversion of a signal into forces for axon outgrowth through Pak1-mediated shootin1 phosphorylation. Current Biology 23, 529–534. .

Trushko, A., Schäffer, E., and Howard, J. (2013). The growth speed of microtubules with XMAP215-coated beads coupled to their ends is increased by tensile force. Proceedings of the National Academy of Sciences 110, 14670–14675. .

Turney, S.G., Ahmed, M., Chandrasekar, I., Wysolmerski, R.B., Goeckeler, Z.M., Rioux, R.M., Whitesides, G.M., and Bridgman, P.C. (2016). Nerve growth factor stimulates axon outgrowth through negative regulation of growth cone actomyosin restraint of microtubule advance. Mol Biol Cell 27, 500–517. .

de Vincentiis, S., Falconieri, A., Mainardi, M., Cappello, V., Scribano, V., Bizzarri, R., Storti, B., Dente, L., Costa, M., and Raffa, V. (2020). Extremely Low Forces Induce Extreme Axon Growth. Journal of Neuroscience https://doi.org/10.1523/JNEUROSCI.3075-19.2020.

Wang, Y., Li, B., Xu, H., Du, S., Liu, T., Ren, J., Zhang, J., Zhang, H., Liu, Y., and Lu, L. (2020). Growth and elongation of axons through mechanical tension mediated by fluorescent-magnetic bifunctional Fe_3_O_4_·Rhodamine 6G@PDA superparticles. Journal of Nanobiotechnology https://doi.org/10.1186/s12951-020-00621-6.

Weiss, P., and Hiscoe, H.B. (1948). Experiments on the mechanism of nerve growth. Journal of Experimental Zoology 107, 315–395. .

Welte, M.A. (2004). Bidirectional Transport along Microtubules. Current Biology 14, R525–R537. https://doi.org/10.1016/j.cub.2004.06.045.

Wen, Q., Po, M.D., Hulme, E., Chen, S., Liu, X., Kwok, S.W., Gershow, M., Leifer, A.M., Butler, V., Fang-Yen, C., Kawano, T., Schafer, W.R., Whitesides, G., Wyart, M., Chklovskii, D.B., Zhen, M., Samuel, A.D.T. (2012). Proprioceptive coupling within motor neurons drives C. elegans forward locomotion Neuron 76, 750–61. doi: 10.1016/j.neuron.2012.08.039.

Yin, J., Straight, P.D., McLoughlin, S.M., Zhou, Z., Lin, A.J., Golan, D.E., Kelleher, N.L., Kolter, R., and Walsh, C.T. (2005). Genetically encoded short peptide tag for versatile protein labeling by Sfp phosphopantetheinyl transferase. Proceedings of the National Academy of Sciences 102, 15815–15820. https://doi.org/10.1073/pnas.0507705102.

Yogev, S., and Shen, K. (2017). Establishing Neuronal Polarity with Environmental and Intrinsic Mechanisms. Neuron 96, 638–650. https://doi.org/10.1016/j.neuron.2017.10.021.

Yogev, S., Cooper, R., Fetter, R., Horowitz, M., and Shen, K. (2016). Microtubule Organization Determines Axonal Transport Dynamics. Neuron 92, 449–460. https://doi.org/10.1016/j.neuron.2016.09.036.

Zhang, W., and Benson, D.L. (2001). Stages of Synapse Development Defined by Dependence on F-Actin. The Journal of Neuroscience 21, 5169–5181. https://doi.org/10.1523/JNEUROSCI.21-14-05169.2001.

Zheng, C., Diaz-Cuadros, M., Nguyen, K.C.Q., Hall, D.H., and Chalfie, M. (2017). Distinct effects of tubulin isotype mutations on neurite growth in *Caenorhabditis elegans*. Molecular Biology of the Cell 28, 2786–2801. https://doi.org/10.1091/mbc.e17-06-0424.

