## Supplementary for "Axonal plasticity in response to active forces generated through magnetic nanopulling"

**Supplemental information**

Table S1. qPCR analysis

| Group | Replicate | Cycle, β-actin mRNA | Cycle, H1 mRNA |
| --- | --- | --- | --- |
| Control, A | 1 | 34.38 | - |
|  | 2 | 38.22 | - |
|  | 3 | 36.77 | - |
| Stretch, A | 1 | 34.87 | - |
|  | 2 | 38.87 | - |
|  | 3 | 38.16 | - |
| Control, S | 1 | 19.97 | 24.92 |
|  | 2 | 19.61 | 24.40 |
|  | 3 | 19.80 | 25.33 |
| Stretch, S | 1 | 19.18 | 24.54 |
|  | 2 | 17.63 | 23.46 |
|  | 3 | 19.68 | 25.230 |

Table S2. GO of cellular Component (CC)

| **GOTERM_cc_direct** | **Term** | **Gene count** | ***P* value** |
| --- | --- | --- | --- |
| GO:0005737 | [cytoplasm](http://www.ebi.ac.uk/QuickGO/GTerm?id=GO:0005737) | 313 | 3.00E-05 |
| GO:0005765 | [lysosomal membrane](http://www.ebi.ac.uk/QuickGO/GTerm?id=GO:0005765) | 24 | 4.90E-05 |
| GO:0005794 | [Golgi apparatus](http://www.ebi.ac.uk/QuickGO/GTerm?id=GO:0005794) | 74 | 8.80E-05 |
| GO:0005768 | [endosome](http://www.ebi.ac.uk/QuickGO/GTerm?id=GO:0005768) | 39 | 4.20E-04 |
| GO:0031902 | [late endosome membrane](http://www.ebi.ac.uk/QuickGO/GTerm?id=GO:0031902) | 12 | 6.30E-04 |
| GO:0005764 | [lysosome](http://www.ebi.ac.uk/QuickGO/GTerm?id=GO:0005764) | 26 | 1.40E-03 |
| GO:0016020 | [membrane](http://www.ebi.ac.uk/QuickGO/GTerm?id=GO:0016020) | 314 | 1.60E-03 |
| GO:0005739 | [mitochondrion](http://www.ebi.ac.uk/QuickGO/GTerm?id=GO:0005739) | 91 | 2.70E-03 |
| GO:0005856 | [cytoskeleton](http://www.ebi.ac.uk/QuickGO/GTerm?id=GO:0005856) | 61 | 7.40E-03 |
| GO:0005815 | [microtubule organizing center](http://www.ebi.ac.uk/QuickGO/GTerm?id=GO:0005815) | 14 | 7.90E-03 |
| GO:0031225 | [anchored component of membrane](http://www.ebi.ac.uk/QuickGO/GTerm?id=GO:0031225) | 13 | 9.40E-03 |
| GO:0005622 | [intracellular](http://www.ebi.ac.uk/QuickGO/GTerm?id=GO:0005622) | 81 | 1.30E-02 |
| GO:0005783 | [endoplasmic reticulum](http://www.ebi.ac.uk/QuickGO/GTerm?id=GO:0005783) | 67 | 2.40E-02 |
| GO:0005770 | [late endosome](http://www.ebi.ac.uk/QuickGO/GTerm?id=GO:0005770) | 11 | 3.30E-02 |
| GO:0045202 | [synapse](http://www.ebi.ac.uk/QuickGO/GTerm?id=GO:0045202) | 28 | 6.40E-02 |
| GO:0005829 | [cytosol](http://www.ebi.ac.uk/QuickGO/GTerm?id=GO:0005829) | 83 | 7.40E-02 |

Table S3. Annotated genes dysregulated in GO:0005856 (cytoskeleton)

| **Symbol** | **Name** | **Up/Down** | **Assigned cathegory** |
| --- | --- | --- | --- |
| Flnb | filamin, beta | up | Actin-binding |
| Mical1 | microtubule associated monooxygenase, calponin and LIM domain containing 1 | up | Actin cytoskeleton |
| Pdlim1 | PDZ and LIM domain 1 (elfin) | up | Actin-binding |
| Svil | supervillin | up | Actin-binding |
| Tln2 | talin 2 | up | Actin-binding |
| Tmod1 | tropomodulin 1 | up | Actin cytoskeleton |
| Tpm2 | tropomyosin 2, beta | up | Actin cytoskeleton |
| Tagln | transgelin | up | Actin cytoskeleton |
| Ahnak | AHNAK nucleoprotein (desmoyokin) | up | Actin cytoskeleton |
| Axl | AXL receptor tyrosine kinase | up | Actin-binding |
| Dgkh | diacylglycerol kinase, eta | up | Actin-binding |
| Trip6 | thyroid hormone receptor interactor 6 | up | Actin-binding |
| Cdc42ep1 | CDC42 effector protein (Rho GTPase binding) 1 | up | Actin cytoskeleton |
| Rgcc | regulator of cell cycle | up | MT cytosckeleton |
| Sh3pxd2b | SH3 and PX domains 2B | up | Actin cytoskeleton |
| Dynlrb2 | dynein light chain roadblock-type 2 | up | Motor protein / transport |
| Rab29 | RAB29, member RAS oncogene family | up | Motor protein / transport |
| Zmynd10 | zinc finger, MYND domain containing 10 | up | Motor protein / transport |
| Kif18b | kinesin family member 18B | up | Motor protein / transport |
| 1110017D15Rik | RIKEN cDNA 1110017D15 gene | up | MT-binding |
| Trim36 | tripartite motif-containing 36 | up | MT-binding |
| Rad51d | RAD51 paralog D | up | MT-binding |
| Ttll9 | tubulin tyrosine ligase-like family, member 9 | up | MT-binding |
| Ccsap | centriole, cilia and spindle associated protein | up | MT-binding |
| Cenpe | centromere protein E | up | MT-binding |
| Fam83d | family with sequence similarity 83, member D | up | MT-binding |
| Mtus1 | mitochondrial tumor suppressor 1 | up | MT-binding |
| Reep4 | receptor accessory protein 4 | up | MT-binding |
| Gramd3 | GRAM domain containing 3 | up | MT cytosckeleton |
| Frmd5 | FERM domain containing 5 | up | MT cytosckeleton |
| Anapc7 | anaphase promoting complex subunit 7 | up | MT-binding |
| Arl6 | ADP-ribosylation factor-like 6 | up | MT-binding |
| Aurkb | aurora kinase B | up | MT cytosckeleton |
| Cenpv | centromere protein V | up | MT-binding |
| Esrra | estrogen related receptor, alpha | up | MT-binding |
| Hmmr | hyaluronan mediated motility receptor (RHAMM) | up | MT-binding |
| Prkci | protein kinase C, iota | up | MT cytosckeleton |
| Abca2 | ATP-binding cassette, sub-family A (ABC1), member 2 | up | MT-binding |
| Pcnt | pericentrin (kendrin) | up | MT-binding |
| Tubgcp5 | tubulin, gamma complex associated protein 5 | up | MT cytosckeleton |
| Bcas3 | BCAS3 microtubule associated cell migration factor | up | MT cytosckeleton |
| Klhl21 | kelch-like 21 | up | MT cytosckeleton |
| Tada2a | transcriptional adaptor 2A | up | MT cytosckeleton |
| Hap1 | huntingtin-associated protein 1 | up | Motor protein / transport |
| Arc | activity regulated cytoskeletal-associated protein | down | Actin cytoskeleton |
| Ccdc66 | coiled-coil domain containing 66 | down | MT-binding |
| Dapk1 | death associated protein kinase 1 | down | Actin cytoskeleton |
| Fntb | farnesyltransferase, CAAX box, beta | down | MT cytosckeleton |
| Gdpd2 | glycerophosphodiester phosphodiesterase domain containing 2 | down | Actin cytoskeleton |
| Katnal1 | katanin p60 subunit A-like 1 | down | MT cytosckeleton |
| Klhl3 | kelch-like 3 | down | Actin-binding |
| Mark4 | MAP/microtubule affinity regulating kinase 4 | down | MT cytosckeleton |
| Mical3 | microtubule associated monooxygenase, calponin and LIM domain containing 3 | down | Actin cytoskeleton |
| Rhobtb3 | Rho-related BTB domain containing 3 | down | Actin cytoskeleton |
| Shtn1 | shootin 1 | down | MT cytosckeleton |
| Spire2 | spire type actin nucleation factor 2 | down | Actin cytoskeleton |
| Topbp1 | topoisomerase (DNA) II binding protein 1 | down | Actin cytoskeleton |
| Tppp | tubulin polymerization promoting protein | down | MT cytosckeleton |
| Tsc1 | TSC complex subunit 1 | down | Actin cytoskeleton |
| Ttll4 | tubulin tyrosine ligase-like family, member 4 | down | MT-binding |

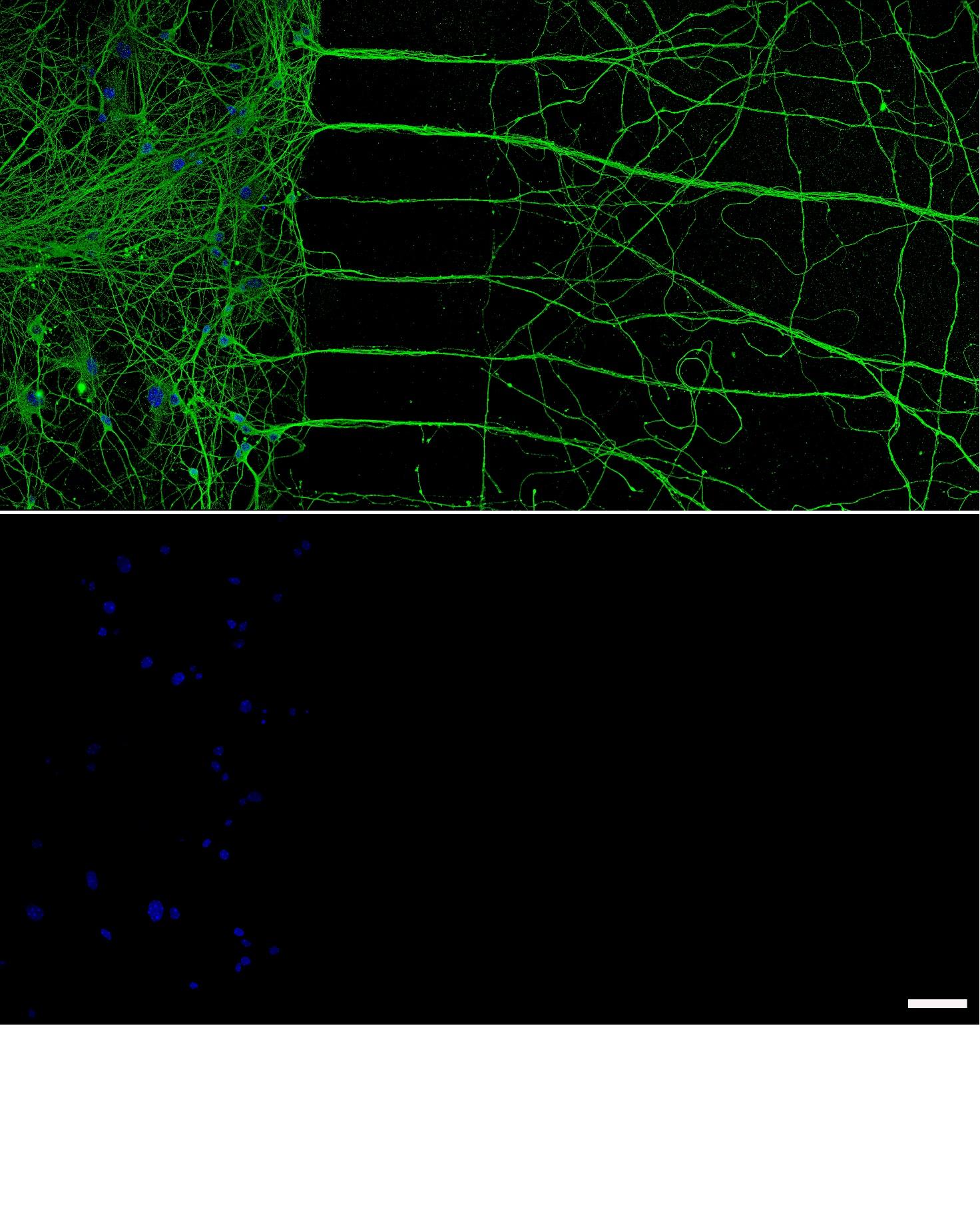

**Figure S1.** Representative image of the somato-dendritic and the axonal compartment. In bottom panel is shown the absence of somata in the axonal compartment as no nuclei were found; BTUBBIII (green), Hoechst (blue) staining; scale bar: 50 μm.

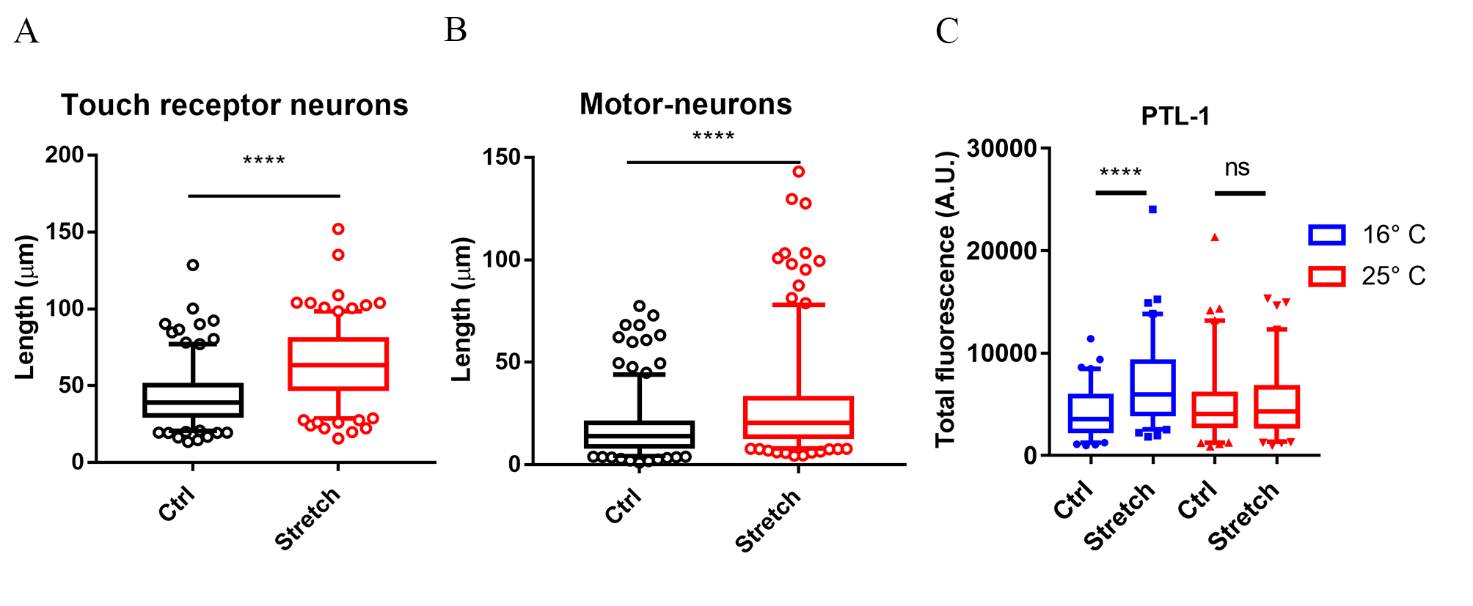

**Figure S2.** (A) Axonal length of motor-neurons in control and stretched conditions (MSB32 strain). Box plot, 9-95 percentile, n=250 from five independent assays. Mann-Whitney test, *p*<0.0001. (B) Axonal length of WT touch receptor neurons for α and β-tubulin (GN692 strain). Control and stretched axons have been measured both, at 16°C and at 25°C. Box plot, min-to-max, n=50 neurites. Kruskal-Wallis test with post hoc Dunn’s test, *p*<0.0001. (C) Quantification of mNG total fluorescence in a transgenic model of tagged mNG::PTL-1 at 16°C and at 25°C. Box plot (5-95 percentile), n=80 axons from four biological replicates Kruskal-Wallis test with post hoc Dunn’s test, *p*<0.0001

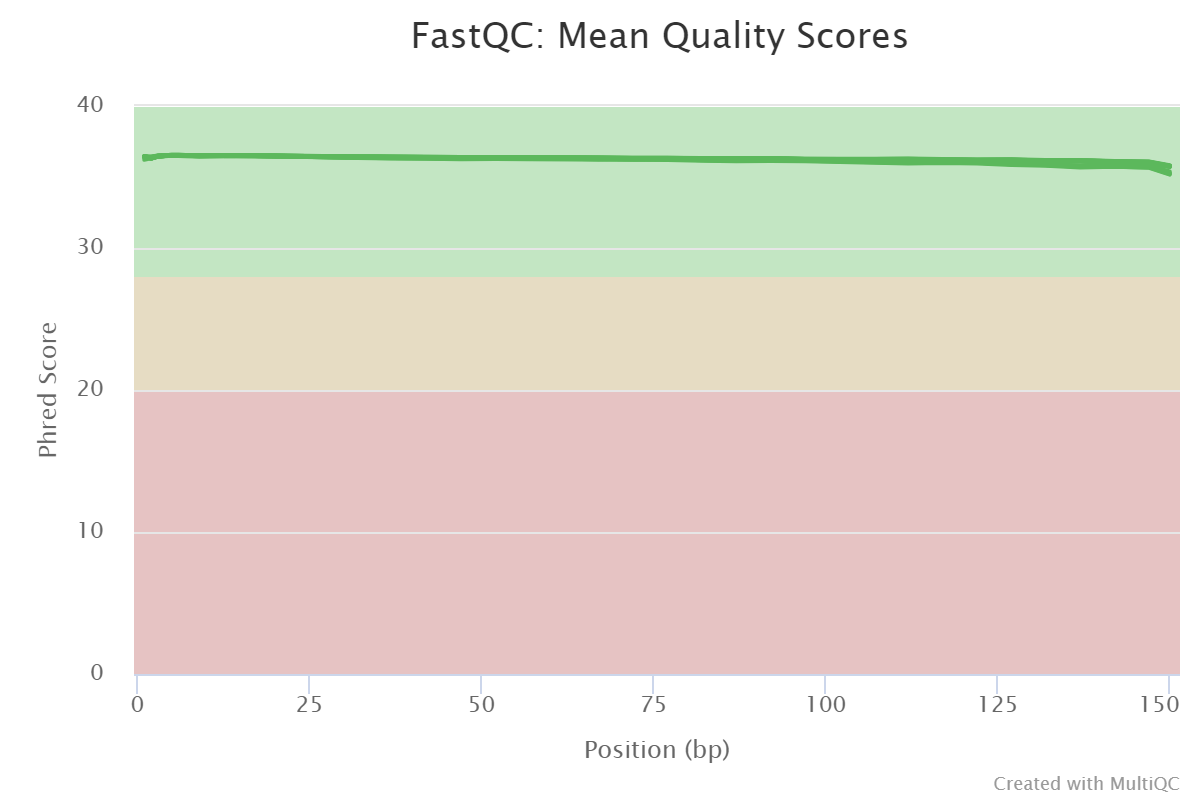

**Figure S3.** Quality control checks on raw sequence datasets.

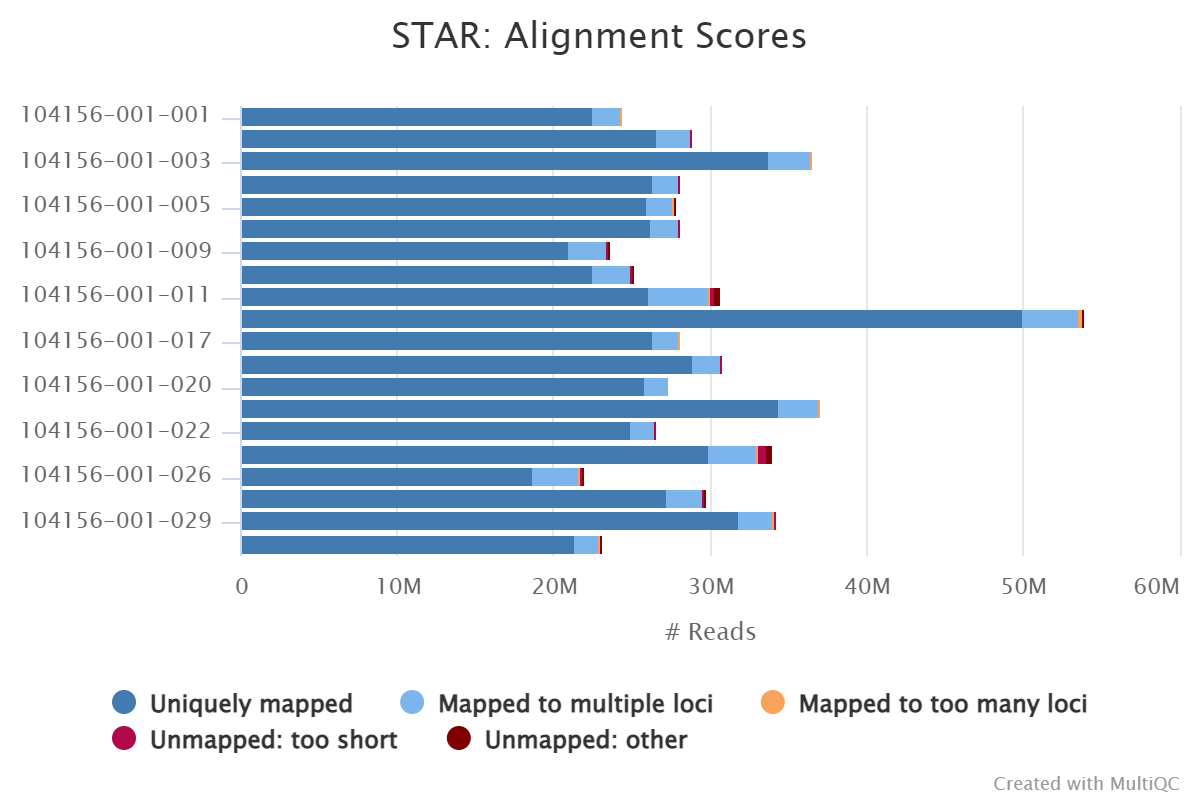

**Figure S4.** Scores of the reads alignment to the reference genome

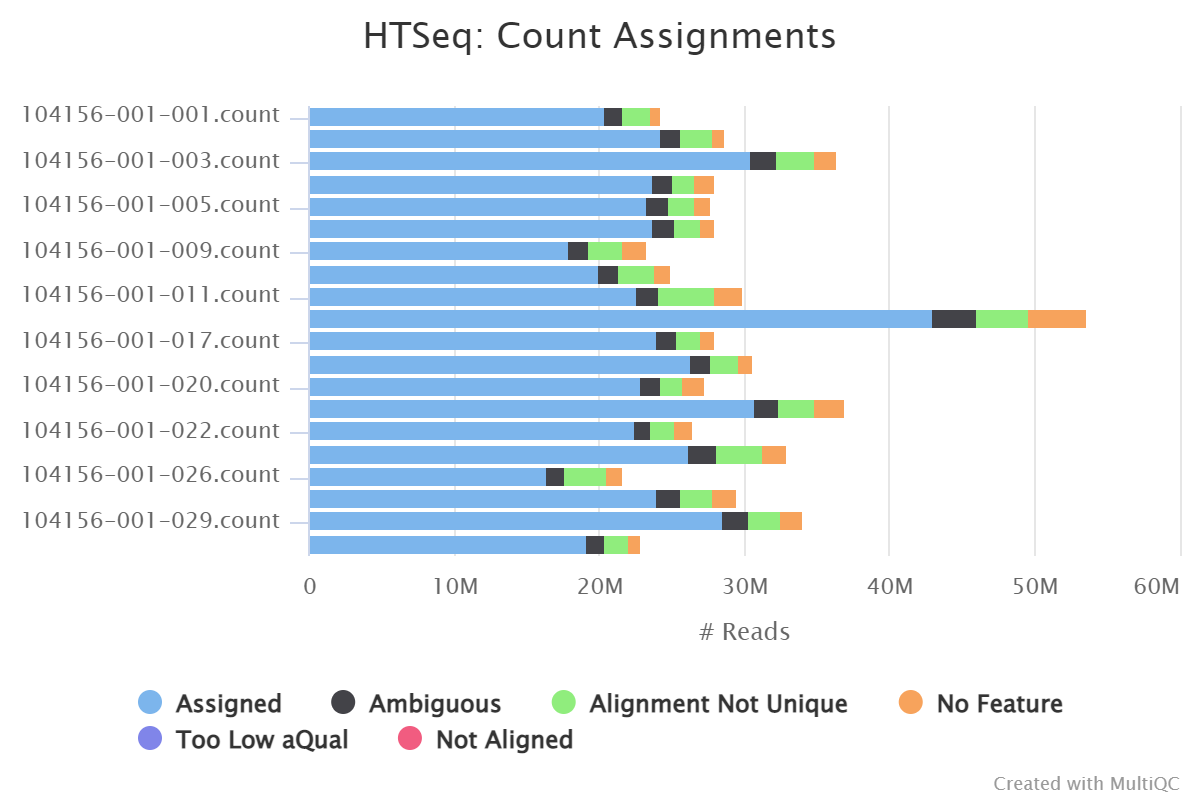

**Figure S5.** Count of the reads mapped against the reference genome.

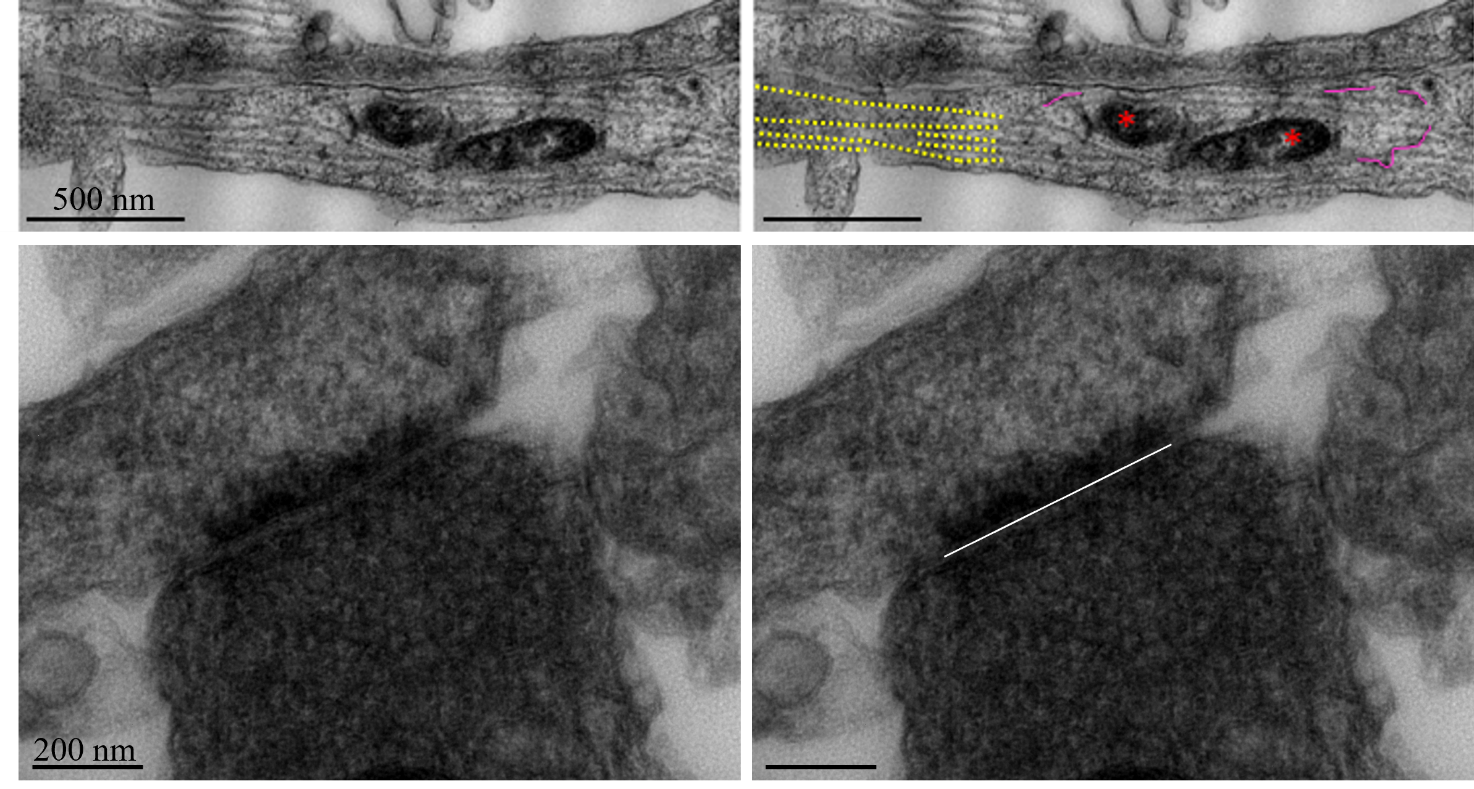

**Figure S6.** Representative TEM images. MTs: dashed yellow lines; ER: magenta lines; mitochondrion indicated with “*”; PSD region: white line.
